# The mechanism of the nucleo-sugar selection by multi-subunit RNA polymerases

**DOI:** 10.1101/2020.06.30.179606

**Authors:** Janne J. Mäkinen, Yeonoh Shin, Eeva Vieras, Pasi Virta, Mikko Metsä-Ketelä, Katsuhiko S. Murakami, Georgiy A. Belogurov

**Author notes:** To whom correspondence should be addressed (K.S.M.) and (G.A.B).

## Abstract

RNA polymerases (RNAPs) synthesize RNA from NTPs, whereas DNA polymerases synthesize DNA from 2’dNTPs. DNA polymerases select against NTPs by using steric gates to exclude the 2’ OH, but RNAPs have to employ alternative selection strategies. In single-subunit RNAPs, a conserved Tyr residue discriminates against 2’dNTPs, whereas selectivity mechanisms of multi-subunit RNAPs remain hitherto unknown. Here we show that a conserved Arg residue uses a two-pronged strategy to select against 2’dNTPs in multi-subunit RNAPs. The conserved Arg interacts with the 2’OH group to promote NTP binding, but selectively inhibits incorporation of 2’dNTPs by interacting with their 3’OH group to favor the catalytically-inert 2’-endo conformation of the deoxyribose moiety. This deformative action is an elegant example of an active selection against a substrate that is a substructure of the correct substrate. Our findings provide important insights into the evolutionary origins of biopolymers and the design of selective inhibitors of viral RNAPs.

## Introduction

All cellular lifeforms use two types of nucleic acids, RNA and DNA to store, propagate and utilize their genetic information. RNA polymerases (RNAPs) synthesize RNA from ribonucleoside triphosphates (NTPs), whereas DNA polymerases (DNAPs) use 2’-deoxyribonucleoside triphosphates (2’dNTPs) to synthesize DNA. The RNA building blocks precede the DNA building blocks biosynthetically and possibly also evolutionarily ^1,2^. Messenger RNA molecules function as information carriers in a single-stranded form, whereas ribosomal, transfer and regulatory RNAs adopt complex three-dimensional structures composed of double-stranded segments. The double stranded RNAs favor A-form geometry where the ribose moiety of each nucleotide adopts the 3’-endo conformation (**Fig. 1a**). In contrast, DNA functions as a B-form double helix, where the deoxyribose of each nucleotide adopts the 2’-endo conformation (**Fig. 1a, b**). Hybrid duplexes between the RNA and DNA transiently form during transcription and adopt an A-form geometry because 2’OH groups in the RNA clash with the phosphate linkages in the B-form configuration. The sugar moieties of NTPs and 2’dNTPs equilibrate freely between the 3’ and 2’-endo conformations in solution with the overall bias typically shifted towards the 2’-endo conformers ^3^. However, both NTPs and 2’dNTPs typically adopt the 3’-endo conformation in the active sites of the nucleic acid polymerases ^4^.

**Fig. 1:**
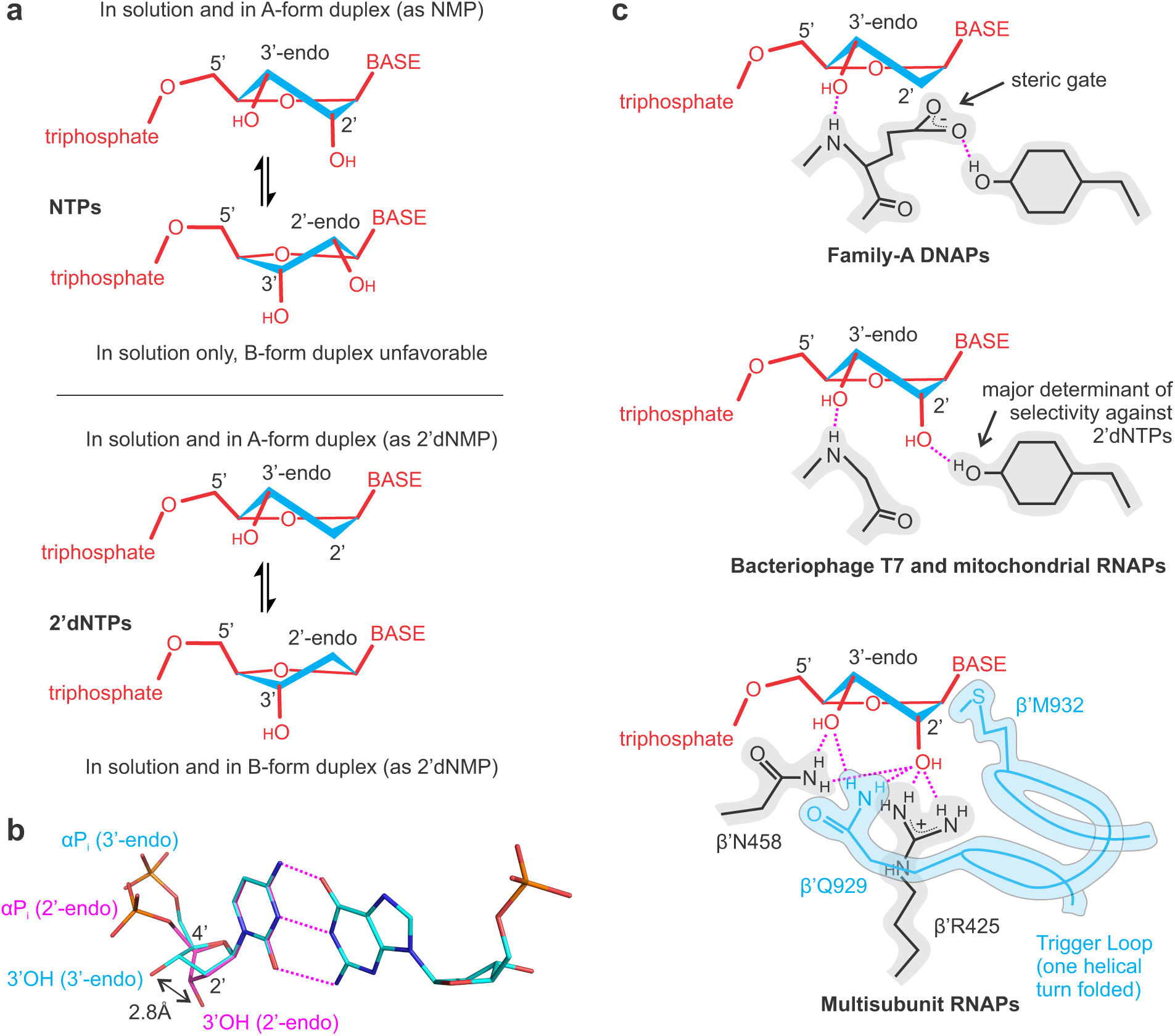
Recognition of NTPs and 2’dNTPs by nucleic acid polymerases. **a** NTPs and 2’dNTPs equilibrate freely between 3’-endo and 2’-endo conformations in solution. A- and B-form duplexes of RNA and DNA constrain sugar moieties of nucleotides in 3’-endo and 2’-endo conformations, respectively. **b** Watson-Crick base pairing of alternative conformers to the template nucleic acid results in markedly different positions of the 3’OH group and the α -phosphate relative to the base-pairing plane. **c** DNAPs and RNAPs stabilize 3’-endo conformers of their cognate substrates. The illustrations reflect the crystallographically documented arrangement of the active site residues but are not accurate projections of 3D structures. Dashed magenta lines depict polar interactions and hydrogen bonds.

RNAPs and DNAPs need to discriminate efficiently against the substrates with the non-cognate sugar. The intracellular levels of NTPs are in the range of hundreds of micromoles to several millimoles per liter and exceed those of the corresponding 2’dNTPs more than tenfold ^5–7^. When selecting the 2’dNTPs, most DNAPs use bulky side-chain residues in their active sites to exclude the 2’OH of the NTPs (reviewed in Ref. ^8^). These steric gate residues, typically Gln/Glu in A-family DNAPs and Tyr/Phe in Y-and B-family DNAPs, create a stacking interaction with the deoxyribose moiety of an incoming 2’dNTP and form a hydrogen bond between the backbone amide group and the 3′-OH group of the deoxyribose moiety (**Fig. 1c**).

Selection against the 2’dNTPs by RNAPs is a daunting challenge because 2’dNTPs are substructures of the corresponding NTPs. Single-subunit RNAPs (e.g., mitochondrial and bacteriophage T7 and N4 enzymes) are homologous and structurally similar to DNAPs. However, single-subunit RNAPs lack a steric gate and use a conserved Tyr residue to discriminate against 2’dNTPs ^9,10^. Tyr selectively facilitates the binding of NTPs by forming a hydrogen bond with the 2’OH group of the NTP ribose (**Fig. 1c**) ^11,12^. Intriguingly, the same Tyr also inhibits the incorporation of 2’dNTPs by an unknown mechanism ^9,13^. Noteworthy, a homologous Tyr hydrogen bonds with the steric gate Gln/Glu residue in A-family DNAPs (**Fig. 1c**)^14,15^.

The mechanism of discrimination against 2’dNTPs by the multi-subunit RNAPs (bacterial, archaeal and eukaryotic nuclear RNAPs) is poorly understood. The combined structural evidence (reviewed in Ref. ^16^) suggests that the 2’OH group can make polar contacts with three universally conserved amino acid side chains: β’Arg425, β’Asn458 and β’Gln929 (numbering of the *Escherichia coli* RNAP). The β’Arg425 and the β’Asn458 are contributed by the active site cavity and can interact with the 2’OH of NTPs in the open and closed active site (see below), whereas the β’Gln929 is contributed by a mobile domain called the trigger loop (TL) and can only transiently interact with the 2’- and 3’-OH of NTPs in the closed active site ^17–19,20^ (**Fig. 1c**).

Closure of the active site by the TL is an essential step during nucleotide incorporation by the multi-subunit RNAPs because the α-phosphate of the NTP is located 5.4 - 6 Å away from the RNA 3’ end in the open active site ^21,22^. Complete closure of the active site by the folding of two alpha-helical turns of the TL positions the triphosphate moiety of the substrate NTP inline for an attack by the 3’OH group of the RNA and accelerates catalysis ∼10^4^ fold ^17,21,23,24^. In contrast, folding of one helical turn of the TL is insufficient to promote catalysis (3’OH → αP distance 5.4 Å ^19^) but likely significantly reduces the rate of NTP dissociation from the active site by establishing contacts between the β’Gln929 and the ribose moiety and stacking of the β’Met932 with the nucleobase (reviewed in Ref. ^16^).

The relative contribution of the TL (β’Gln929 and β’Met932) and the active site cavity (β’Arg425 and β’Asn458) to the discrimination against 2’dNTPs remains hitherto uncertain. The closure of the active site makes only a 5- to 10-fold contribution to an overall 500- to 5000-fold selectivity against the 2’dNTP in RNAPs from *E. coli* ^24^ and *Saccharomyces cerevisiae* ^25^. Consistently, the open active site of the *E. coli* RNAP retained a ∼100-fold overall selectivity against 2’dNTPs ^24^. However, the open active site of the *Thermus aquaticus* RNAP has been reported to be largely unselective ^23^, and individual substitutions of the β’Asn458 with Ser in *E. coli* and *S. cerevisiae* resulted in only a <5-fold decrease in selectivity ^17,26^. Most importantly, although the universally conserved β′Arg425 closely approaches 2’OH of the NTP in several X-ray crystal structures ^17–19,22,20^ (**Supplementary Table 5**) and has been highlighted as the sole residue mediating the selectivity against 2’dNTP in a computational study by Roßbach and Ochsenfeld ^27^, the role of this residue has not been experimentally assessed.

In this study, we systematically investigated the effects of individual substitutions of the active site residues on the discrimination against 2’dNTPs in single nucleotide addition (SNA) assays and during processive transcript elongation by the *E. coli* RNAP. This analysis demonstrated that β’Arg425 is the major determinant of the selectivity against 2’dNTPs in multi-subunit RNAPs. We further analyzed the binding of 2’-deoxy substrates by *in silico* docking and X-ray crystallography of *Thermus thermophilus* RNAP. Our data suggest that the conserved Arg actively selects against 2’dNTPs by favoring their templated binding in the 2’-endo conformation that is poorly suitable for incorporation into RNA.

## Results

### β’Arg425 is the major determinant of the selectivity against the 2’dGTP by *E. coli* RNAP

To investigate the mechanism of the discrimination against the 2’-deoxy substrates we performed time-resolved studied of the single nucleotide incorporation by the wild-type (WT) and variant *E. coli* RNAPs. Among several single substitutions of the key residues that contact NTP ribose (**Fig. 1c**), we selected four variant RNAPs that retained at least half of the wild-type activity at saturating concentration of NTPs. This approach minimized the possibility that the amino acid substitutions induced global rearrangements of the active site thereby complicating the interpretations of their effects on the sugar selectivity.

Transcript elongation complexes (TECs) were assembled on synthetic nucleic acid scaffolds and they contained the fully complementary transcription bubble flanked by 20-nucleotide DNA duplexes upstream and downstream (**Supplementary Fig. 1a**). The annealing region of a 16-nucleotide RNA primer was initially 9 nucleotides, permitting the TEC extended by one nucleotide to adopt the post- and pre-translocated states, but disfavoring backtracking ^28^. The RNA primer was 5’ labeled with the infrared fluorophore ATTO680 to monitor the RNA extension by denaturing PAGE. The template DNA strand contained the fluorescent base analogue 6-methyl-isoxanthopterin (6-MI) eight nucleotides upstream from the RNA 3’ end to monitor RNAP translocation along the DNA following nucleotide incorporation ^29^.

We first measured GTP and 2’dGTP concentration series of the WT and altered RNAPs using a time-resolved fluorescence assay performed in a stopped-flow instrument (**Supplementary Fig. 1b, c**). We used the translocation assay because it allowed rapid acquisition of concentration series, whereas measurements of concentration series by monitoring RNA extension in the rapid chemical quench-flow setup would be considerably more laborious. The concentration series data allowed the estimation of *k*_*cat*_ and the *Km* (Michaelis constant) for GTP and 2’dGTP. We then supplemented the concentration series with timecourses of GMP and 2’dGMP incorporation obtained using a rapid chemical quench-flow technique with EDTA as a quencher. EDTA inactivates the free GTP and 2’dGTP by chelating Mg^2+^ but allows a fraction of the bound substrate to complete incorporation into RNA after the addition of EDTA ^30,31^. As a result, the EDTA quench experiment is equivalent to a pulse-chase setup and provides information about the rate of substrate dissociation from the active site of RNAP. A global analysis of the concentration series and EDTA quench experiments (*i*) allowed the estimation of the *K*_*D*_ for GTP dissociation from the active site and (*ii*) suggested that the *K*_*D*_ for the dissociation of the 2’dGTP from the active site approximately equals the *Km* for 2’dGMP incorporation (see **Supplementary Note**). We further used inferred values of *k*_*cat*_ and *K*_*D*_ to compare the capabilities of the variant RNAPs to discriminate against 2’dGTP (**Fig. 2**).

**Fig. 2:**
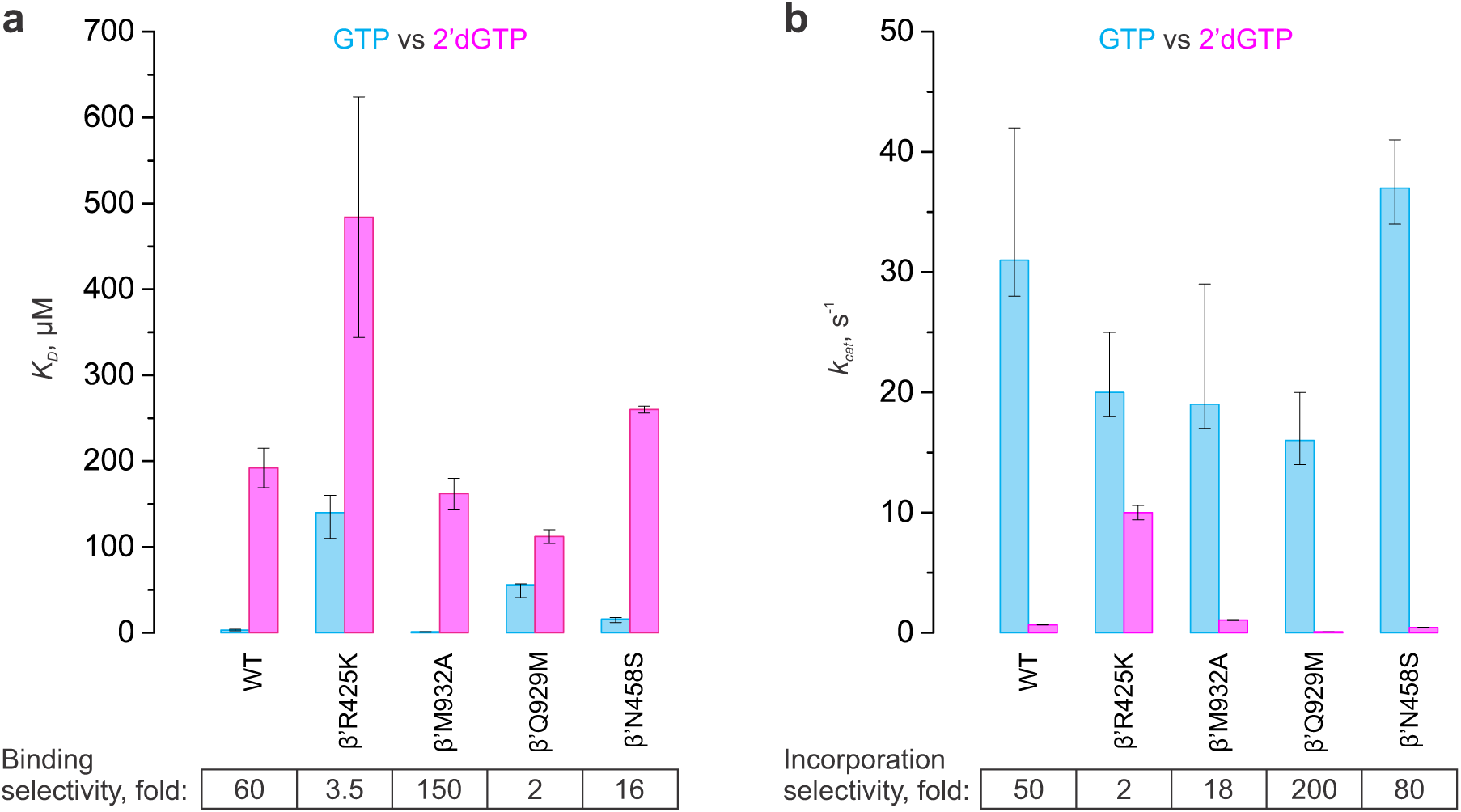
Nucleo-sugar selectivities of the WT and variant *E. coli* RNAP. **a** Equilibrium dissociation constants for the reversible binding of GTP and 2’dGTP. **b** Turnover numbers for the incorporation of GMP and 2’dGMP. Numerical values of the parameters are indicated in **Table 1** and **Supplementary Tables 1-2**. Error bars are ranges of duplicate measurements or SDs of the best-fit parameters, whichever values were larger.

WT RNAP displayed ∼60-fold higher affinity for GTP than for 2’dGTP (**Fig. 2a, Table 1**). The β’R425K and β’Q929M substitutions decreased the selectivity at the binding step 17- and 30- fold, respectively, largely by decreasing the affinity for GTP. In contrast, the β’N458S decreased the selectivity only 4-fold, whereas the β’M932A increased the selectivity 2.5-fold. At saturating substrate concentrations, the WT RNAP incorporated GMP ∼50-fold faster than the 2’dGMP (**Fig. 2b, Table 1**). The β’R425K substitution decreased the selectivity 25-fold, primarily by accelerating the incorporation of the 2’dGMP. In comparison, the effects of other substitutions on the selectivity against the 2’dGTP at the incorporation step were relatively small (**Fig. 2b, Supplementary Tables 3, 4**). The β’M932A decreased the selectivity 3-fold, whereas the β’N458S and β’Q929M increased the selectivity 1.5- and 4-fold, respectively. Noteworthy, the β’Q929M decreased the rate of 2’dGMP incorporation 10-fold.

**Table 1.** Binding and incorporation of the GMP, 2’dGMP and 3’dGMP by the wild-type and β’R425K *E. coli* RNAPs.

| **Scheme 1:** $\text{TEC16} + \text{GTP} \xrightleftharpoons[k_{\text{off}}]{k_{\text{on}}} \text{TEC16GTP} \xrightarrow{k_{\text{cat}}} \text{TEC17}_{\text{PRE}} \xrightarrow{k_{\text{trn}}} \text{TEC17}_{\text{POST}}$ |  |  |  |  |  |  |  |
| --- | --- | --- | --- | --- | --- | --- | --- |
| RNAP | Substrate | $k_{\text{on}}$, $\mu\text{M}^{-1}\text{s}^{-1}$ | $k_{\text{off}}$, $\text{s}^{-1}$ | $k_{\text{cat}}$, $\text{s}^{-1}$ (fast) | $k_{\text{cat}}$, $\text{s}^{-1}$ (slow) | fast fraction, % | $K_D$, $\mu\text{M}$ |
| WT | GTP | 2.5 (2.2 – 3.0) | 7.7 (4.0 – 9.6) | 31 (28 – 42) | 2.6 (0.8 – 7.6) | 88 (77 – 93) | 3.1 (2.3 – 4.0) |
| WT | 2'dGTP | > 0.24 | > 45 | 1.4 (1.1 – 2.3) | 0.35 (0.22 – 0.49) | 52 (30 – 70) | 190 (160 – 220) |
| $\beta$'R425K | GTP | 0.5 (0.4 – 0.8) | 72 (57 – 120) | 20 (18 – 25) | 1.3 (0.4 – 4.1) | 86 (76 – 92) | 140 (110 – 160) |
| $\beta$'R425K | 2'dGTP | > 0.10 | > 30 | 13 (11 – 18) | 2.2 (0.7 – 5.4) | 78 (50 – 90) | 420 (330 – 480) |
| WT | 3'dGTP | 2.7 (2.2 – 4.4) | 87 (70 – 140) | 12 (11 – 13) | 0.74 (0.5 – 1.4) | 80 (75 – 85) | 32 (28 – 38) |

The reaction products were modelled as sums of independent contributions by the fast and slow fractions of RNAP using numerical integration capabilities of the KinTek Explorer software. Contributions of each fraction were modeled as **Scheme 1**. Upper and lower bounds of the parameters were calculated at a 10% increase in $\text{Chi}^2$ .

Overall, these experiments suggested that the β’Arg425 plays a central role in the discrimination against 2’-deoxy substrates (**Table 1**): the β’Arg425 selectively facilitated binding of GTP and selectively inhibited the incorporation of 2’dGMP. In contrast, the role of the β’Gln929 was complex: while the β’Gln929 selectively facilitated the binding of GTP, it also selectively facilitated the incorporation of 2’dGMP.

### β’Arg425 inhibits the utilization of 2’dNTPs during processive transcript elongation

The time-resolved SNA assays described above are superior to any other currently available techniques for the quantitative assessment of the binding and incorporation of different substrates and the effects of active site residues therein. However, these assays have several limitations: the nucleotide incorporation was measured for static complexes stabilized in the post-translocated state by the artificially limited RNA:DNA complementarity and the effects are assessed only at a single, easy to transcribe, sequence position. To test if the conclusions drawn from the SNA assay remain valid during processive transcript elongation we developed a semiquantitative assay as follows.

TECs were assembled on a nucleic acid scaffold with a 49 bp-long downstream DNA and chased with NTP mixtures containing 50 µM ATP, CTP, UTP and GTP or 2’dGTP for 2 min at 25°C. Transcription with the 2’dGTP by the WT RNAP resulted in characteristic pauses at each sequence position preceding the incorporation of the 2’dGMP (**Fig. 3**, *pre-G sites*). We used the amplitude of these accumulations as a semi-quantitative measure of the ability of the RNAP to utilize 2’dGTP. Noteworthy, the interpretation of the processive transcription by some variant RNAPs was complicated by enhanced pausing after the incorporation of cytosine (**Fig. 3b**, *at-C sites*) and 2’dGMP (**Fig. 3b**, *at-G sites*) in certain sequence contexts. However, these additional pauses were unrelated to the utilization of the 2’dGTP as a substrate and could be disregarded when comparing pre-G pauses that occurred upstream of all at-C and at-G pauses.

**Fig. 3:**
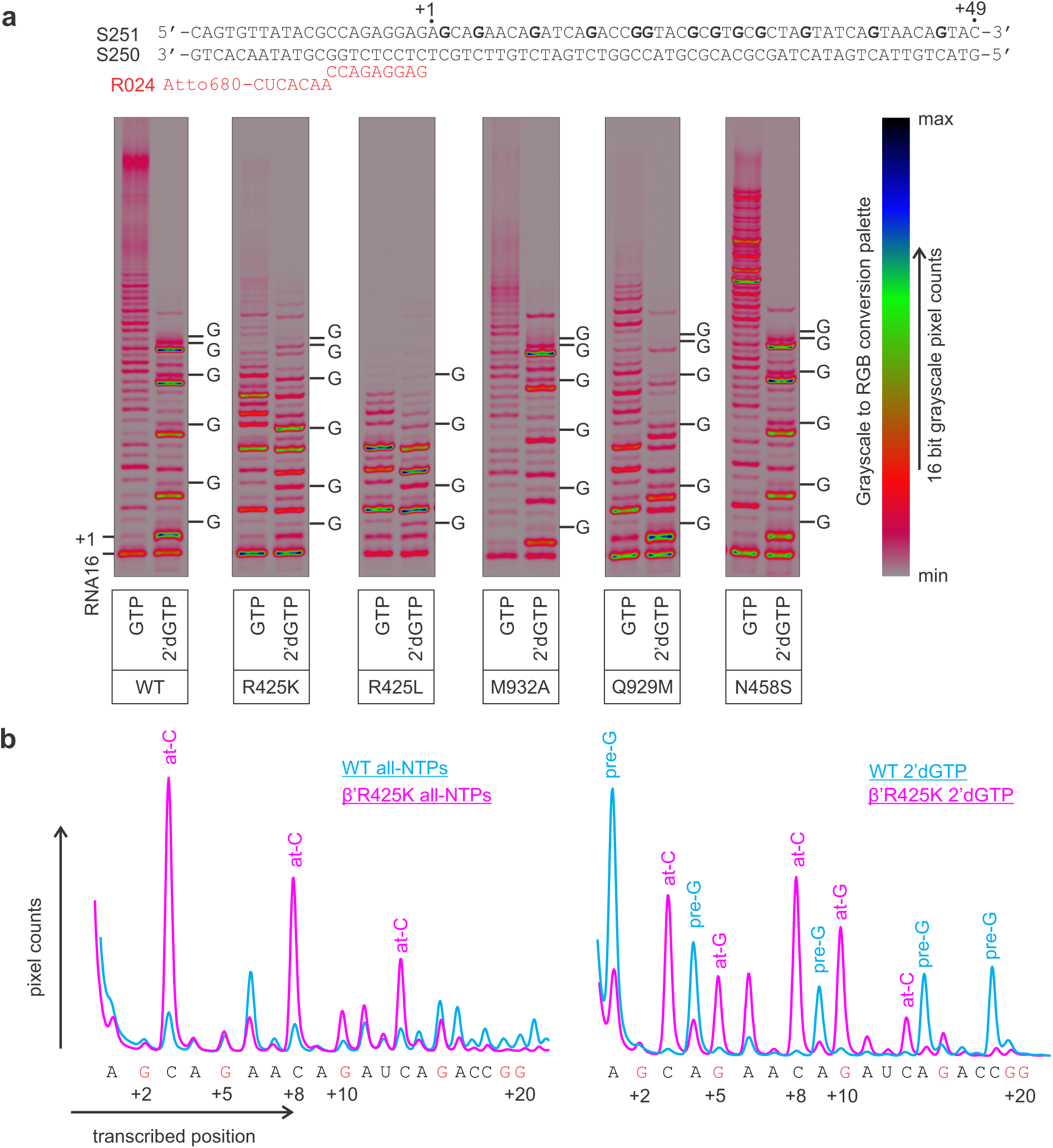
Utilization of 2’ dGTP during the processive transcription by the WT and variant *E. coli* RNAPs. **a** TECs were assembled using the scaffold shown above the gel panels and chased with 50 µM ATP, CTP, UTP and GTP or 2’dGTP for 2 min at 25°C. The positions of GMPs in the resolved stretches of the transcribed sequence are marked along the right edge of the gel panels. 16-bit grayscale scans were normalized using max pixel counts within each gel panel and pseudo-colored using RGB palette on the right. **b** Lane profiles of transcription in all-NTPs and 2’dGTP chases by the wild-type and β’R425K RNAPs quantified from gels in **(a)**. Traces were manually aligned along the X-axis and scaled along the Y-axis using several sequence positions as references.

In contrast to the WT RNAP, the β’R425K did not pause prior to the incorporation of the 2’dGMP (**Fig. 3**), consistently with the significantly higher 2’dGMP incorporation rate observed in SNA assays (**Fig. 2b**). Moreover, the β’R425L RNAP also did not accumulate at the pre-G sites despite being strongly defective during processive transcription (**Fig. 3a, Supplementary Fig. 4**). These data suggest that the loss of selectivity in the β’R425K is attributable to the absence of the β’R425 rather than the gain of function effect of the Lys residue at the corresponding position.

The β’M932A paused noticeably less whereas the β’Q929M paused noticeably more than the WT RNAP at the pre-G sites (**Fig. 3a, Supplementary Fig. 4**) consistently with the 2-fold higher (β’M932A) and 10-fold lower (β’Q932M) *k*_*cat*_ for the 2’dGMP incorporation in the SNA experiments (**Fig. 2b, Table 1**). In contrast, the β’N458S was largely indistinguishable from the WT RNAP in its ability to utilize the 2’dGTP in the processive transcription assay (**Fig. 3, Supplementary Fig. 4**), presumably because this assay is not sensitive enough to resolve the ∼1.5-fold difference in *k*_*cat*_ for the 2’dGMP incorporation (**Fig. 2b, Supplementary Tables 3, 4**). Overall, the analysis of the utilization of the 2’dGTP during the processive transcription of diverse sequences by the WT and variant RNAPs recapitulated the major effects observed in the SNA experiments.

Next, we tested the effects of the β’R425K, β’M932A, β’Q932M and β’N458S substitutions on utilization of 2’dATP, 2’dCTP and 2’dUTP during processive transcription (**Supplementary Figs. 5-7**). For each 2’dNTP, we custom designed a template where the 2’dNTP is incorporated several times early in transcription, thereby allowing unambiguous interpretation of the accumulation of RNAPs at sites preceding the 2’dNMP incorporation. An analysis of the utilization of 2’dATP, 2’dCTP and 2’dUTP largely recapitulated the effects observed for 2’dGTP, except that the β’N458S was markedly inferior to the WT RNAP in utilizing 2’dATP and 2’dUTP. Overall, these data demonstrated that the enhanced or diminished capabilities of the variant RNAPs to utilize 2’dGTP in the SNA assays reflected, in qualitative terms, their capabilities to utilize all four 2’dNTPs.

### β’Arg425 favors the binding of the 2’dCTP in the 2’-endo conformation

The role of the β’Arg425 in selectively promoting the binding of NTPs was easy to explain because the β’Arg425 interacts with the 2’OH of the NTP analogues in several RNAP structures (**Supplementary Table 5, Fig. 1c, 4a, b**). In contrast, the observation that the β’Arg425 selectively inhibited the incorporation of 2’dNTPs could not be readily explained: our results show that the β’Arg425 substitutions promote the incorporation of the substrate that lacks the 2’OH group, which the β’Arg425 would interact with. We hypothesized that, in the absence of the 2’OH, the β’Arg425 interacted with something else and that the interaction slowed down the incorporation of 2’dNMPs into the nascent RNA. We further reasoned that the 3’OH group of the 2’dNTP was the most likely interacting partner of the β’Arg425, an inference supported by MD simulations of *S. cerevisiae* RNAPII ^27^. However, the 3’OH group is positioned too far from the β’Arg425 when the sugar moiety is in the 3’-endo conformation (**Supplementary Table 5**). We further hypothesized and demonstrated by *in silico* docking experiments that the 3’OH could move to within the hydrogen bond distance of the β’Arg425 if the deoxyribose moiety adopted a 2’-endo conformation (**Supplementary Fig. 8, Supplementary Table 6, Supplementary Data 1-3**).

To test this hypothesis *in crystallo*, we solved the X-ray crystal structure of the initially transcribing complexes containing *T. thermophilus* RNAP, DNA and 3-nt RNA primer with incoming 2’dCTP bound at the active site at 3.14 Å resolution. The structure displayed a well-resolved electron density of the 2’dCTP and the β’Arg425 closely approaching the deoxyribose moiety (**Fig. 4c, d, Table 2, Supplementary Fig. 9a, Supplementary Data 4**). The 2’dCTP was observed in the pre-insertion conformation, that was unsuitable for catalysis because the α-phosphate was located 5.7 Å from the 3’OH of the RNA primer. The electron density was consistent with the interaction between the β’Arg425 and the 3’OH group of the deoxyribose in the 2’-endo conformation in agreement with the results of *in silico* docking. Interestingly, the density for the metal ion complexed by the β- and γ-phosphates of the 2’dCTP was weak and the coordination distances were longer than typically observed for Mg^2+^ in the corresponding position. We modeled this metal ion as a Na^+^ rather than Mg^2+^ similarly to what has been proposed for DNA polymerase β ^32^.

**Table 2:** Data collection and refinement statistics

| Complex | 2'dCTP | 3'dCTP |
| --- | --- | --- |
| PDB code | 6WOX | 6WOY |
| **Data collection** |  |  |
| Space group | C2 | C2 |
| Cell dimensions |  |  |
| *a* (Å) | 185.44 | 185.98 |
| *b* (Å) | 102.65 | 102.47 |
| *c* (Å) | 295.38 | 296.04 |
| $\beta$ (°) | 98.91 | 98.87 |
| Resolution (Å) | 30 – 3.14 | 30 – 3.00 |
| Total reflections | 275,994 | 361,353 |
| Unique reflections | 89,931 | 106,804 |
| Redundancy | 3.1 (2.5) | 3.4 (2.9) |
| Completeness (%) | 94.3 (68.6) | 96.9 (73.6) |
| *I* / $\sigma$ | 10.06 (1.44) | 14.78 (1.73) |
| *R*<sub>merge</sub> | 0.129 (0.521) | 0.086 (0.553) |
| CC<sup>1/2</sup> | 0.985 (0.829) | 0.991 (0.855) |
| **Refinement** |  |  |
| Resolution (Å) | 30 – 3.14 | 30 – 3.00 |
| *R*<sub>work</sub> | 0.196 | 0.206 |
| *R*<sub>free</sub> | 0.261 | 0.219 |
| No. of atoms | 28,569 | 28,569 |
| R.m.s deviations |  |  |
| Bond length (Å) | 0.010 | 0.009 |
| Bond angles (°) | 1.22 | 1.12 |
| Clashscore | 15.04 | 13.19 |
| Ramachandran favored, % | 85.68 | 86.95 |
| Ramachandran outliers, % | 0.37 | 0.37 |

Data sets were collected at MacCHESS F1 line, Ithaca, NY
\*Highest resolution shells are shown in parentheses

**Fig. 4:**
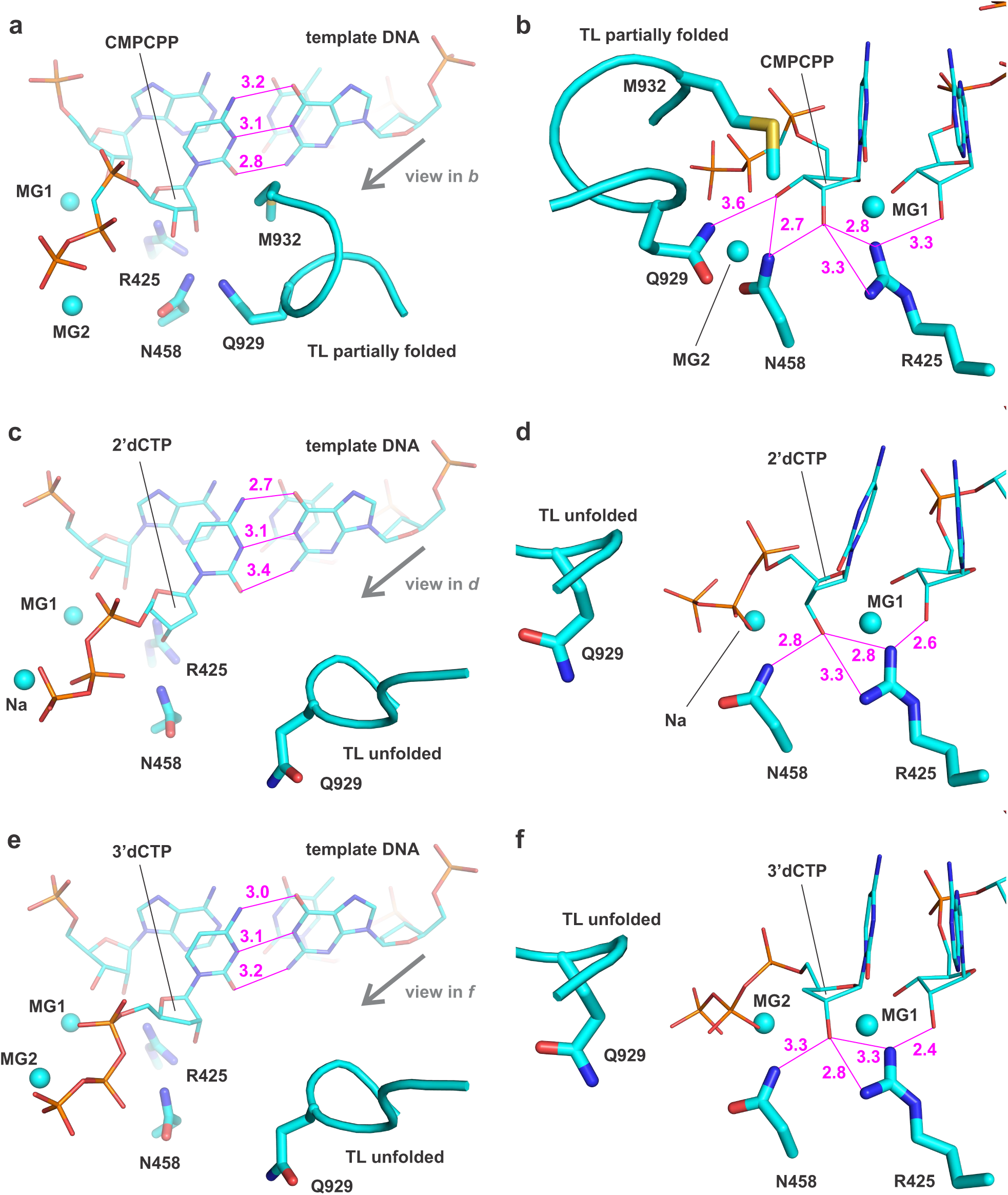
Binding poses of CMPCPP (a, b), 2’dCTP (c, d) and 3’dCTP (e, f) in the active site of *T. thermophilus* RNAP. Magenta numbers are interatomic distances in Å. Panels **(a)** and **(b)** were prepared using PDB ID 4Q4Z ^19^.

The TL was completely unfolded in the structure of the initially transcribing complex with the 2’dCTP, in contrast to a partially helical conformation typically observed in the structures of the RNAP complexes with non-hydrolysable NTP analogues (**Supplementary Table 5, Supplementary Data 5**) ^17–19,20^. The destabilization of the TL folding could be due to the absence of interactions between the β’Gln929 and the 3’OH of the 2’dCTP. The latter effect could in turn be attributed to the sequestration of the 3’OH by the β’Arg425 and the different conformation of the deoxyribose moiety.

To test if the unavailability of the 3’OH group was indeed responsible for the destabilization of the TL folding, we solved the X-ray crystal structure of the initially transcribing complex of the *T. thermophilus* RNAP with a 3’dCTP at 3.0 Å resolution. The structure displayed a well-resolved density of the 3’dCTP and the β’Arg425 closely approaching the 3’-deoxyribose moiety (**Fig. 4e, f, Supplementary Fig. 9b, Supplementary Data 6**). The 3’dCTP was in the pre-insertion conformation that was unsuitable for catalysis because the α-phosphate was located 5.6 Å away from the 3’OH of the RNA primer. The overall pose of the 3’dCTP was similar to that of cytidine-5’-[(α,β)-methyleno]triphosphate (CMPCPP): the 3’-deoxyribose adopted a 3’-endo conformation and the 2’OH group interacted with the β’Arg425. However, the TL was completely unfolded, supporting our hypothesis that the unavailability of the 3’OH group was alone sufficient to significantly destabilize the folding of the first helical turn of the TL.

Overall, the comparative analysis of RNAP structures with CMPCPP, 2’dCTP and 3’dCTP suggested that the β’Arg425 inhibited the incorporation of 2’dNTPs by interacting with their 3’OH group and favoring the 2’-endo conformation of the deoxyribose moiety. At the same time, the structures did not provide a decisive answer as to why the 2’-endo conformations of 2’dNTPs were less suitable for incorporation into RNA than the 3’-endo conformations.

### The misplacement of the 3’OH only partially accounts for the inertness of 2’-endo 2’dNTPs

The X-ray structures and *in silico* modeling experiments suggested that interactions between the 3’OH of the deoxyribose moiety and the β’Arg425 or the β’Gln929 were mutually exclusive. Accordingly, the β’Arg425 could inhibit the incorporation of the 2’dNMP solely by slowing down the initial steps of the TL folding, by sequestering the 3’OH group and preventing its interaction with the β’Gln929 of the TL. To test this hypothesis, we determined the incorporation rate of 3’dGMP by the WT RNAP (**Supplementary Fig. 3c**). We found that the *k*_*cat*_ for 3’dGMP incorporation was only 5-fold slower than the *k*_*cat*_ for GMP incorporation and 10-fold higher than the *k*_*cat*_ for 2’dGMP incorporation (**Table 1**). These data demonstrated that the sequestration of the 3’OH group accounted for no more than a 5-fold inhibition of the 2’dGMP incorporation by the β’Arg425. The remaining 10-fold inhibition of the overall 50-fold inhibitory effect was contributed by some other features of the 2’-endo binding pose, as discussed below.

## Discussion

In this study we performed a systematic analysis of the role of the amino acid residues in the active site of the multi-subunit RNAP in selecting NTPs over 2’dNTPs. We identified a conserved Arg residue, β’Arg425 (*E. coli* RNAP numbering) as the major determinant of the sugar selectivity. The β’Arg425 favored binding of GTP over 2’dGTP and selectively inhibited the incorporation of 2’dNMPs into RNA (**Figs. 2, 3, Table 1**).

The enhancement of NTP binding by the β’Arg425 is consistent with the observation that the β’Arg425 is positioned to hydrogen bond with the 2’OH of the NTP substrate analogues in several RNAP structures (**Supplementary Table 5**) and with MD simulations of the *S. cerevisiae* RNAPII ^27^. However, the existing data fail to explain the inhibition of the 2’dNTP incorporation by the β’Arg425. In search of an explanation, we performed *in silico* docking experiments and solved the X-ray crystal structures of the initially transcribing *T. thermophilus* RNAP with the cognate 2’dCTP and 3’dCTP. These experiments revealed that the β’Arg425 interacts with the 3’OH group of the 2’dNTP substrate and favors the 2’-endo conformation of the deoxyribose (**Fig. 4c d, Supplementary Fig. 8e, f**). In contrast, the ribose of the cognate NTP substrate is stabilized in the 3’-endo conformation by multiple polar contacts and hydrogen bonds with the active site residues: β’Arg425, β’Asn458 and β’Gln929 (**Figs. 1c, 4a, b, Supplementary Fig. 8a, b, Supplementary Table 5**).

We next considered whether the deformation of the 2’dNTP substrate, repositioning of the β’Arg425 or both were behind the slow incorporation of the 2’dNMPs by the WT RNAP. A hybrid quantum and molecular mechanics (QM/MM) analysis of nucleotide incorporation by the *S. cerevisiae* RNAPII suggested that repositioning of the β’Arg425 by the 2’dNTP substrate may increase the activation energy barrier for the nucleotide addition reaction ^27^. However, a comparison of the RNAP structures with bound CMPCPP, 2’dCTP and 3’dCTP revealed very small changes in the conformation of the β’Arg425 (**Fig. 4**). Similarly, a survey of the published X-ray and CryoEM structures revealed that the β’Arg425 occupies approximately the same volume irrespective of the presence or absence of the active site ligands (**Supplementary Table 5)**. Accordingly, we reasoned that the preferential selection of the catalytically-inert 2’-endo conformers of 2’dNTPs and the deformation of the catalytically-labile 3’-endo conformers of 2’dNTPs by β’Arg425 were likely the major factors behind the slow incorporation of the 2’dNMPs. However, it remained unclear why the 2’-endo conformers of the substrates were less suitable for the incorporation than the 3’-endo conformers.

We first explored the possibility that the sequestration of the 3’OH group by the β’Arg425 makes it unavailable for the interaction with the β’Gln929 of the TL (**Fig. 4c, d, Supplementary Fig. 8e, f**), thereby destabilizing the TL-mediated closure of the active site. It is well established that the closure of the active site by two helical turns of the TL accelerates the catalysis of nucleotide incorporation by ∼10^4^-fold ^17,21,23,24^. Noteworthy, the TL is partially folded in most structures with ribonucleotide substrate analogues (**Fig. 4a, b, Supplementary Table 5)** ^17–19,20^, yet was completely unfolded in the structures we obtained with either a 2’dCTP (**Fig. 4c, d**) or a 3’dCTP (**Fig. 4e, f**). Given that the 3’dCTP was in the conventional 3’-endo conformation, the latter result suggested that the unavailability of the 3’OH group was sufficient to significantly impair the folding of the TL and slow down the catalysis by the *T. thermophilus* RNAP *in crystallo*.

To quantitatively estimate the contribution of the 3’OH interactions to the catalysis, we determined the rate of the 3’dGMP incorporation by *E. coli* RNAP. We found that the rate of the 3’dGMP incorporation was 5-fold slower than the rate of the GMP incorporation, but 10-fold faster than that of the 2’dGMP (**Table 1, Supplementary Fig. 3c**). These results suggested that the sequestration of the 3’OH group by the β’Arg425 could account for no more than a 5-fold out of its 50-fold overall inhibitory effect. Notably, the *T. thermophilus* RNAP also incorporates 3’dNMPs faster than 2’dNMPs but discriminates against both types of substrates ∼40-fold stronger than the *E. coli* RNAP ^23^.

Similarly, the effects of the β’Q929M substitution were inconsistent with the idea that the 3’OH capture by β’Arg425 could alone account for the slow rate of the 2’dNMP incorporation. If that were true, the β’Q929M variant should be relatively insensitive to the absence of the 2’OH group. However, the opposite was true: β’Q929M was only twofold slower in incorporating GMP than the WT RNAP, but tenfold slower in incorporating 2’dGMP. We propose that the β’Gln929 competes with the β’Arg425 for the 3’OH group of the 2’dNTP substrate: the β’Arg425 favors the catalytically-inert 2’-endo conformer (**Fig. 4c, d, Supplementary Fig. 8e, f**), whereas the β’Gln929 favors the catalytically-labile 3’-endo conformer (**Supplementary Fig. 8c, d**). As a result, the β’Gln929 is more important during the incorporation of 2’dNMPs than NMPs.

Since the TL folding can account only for a fraction of the inhibitory effect, what other factors make the 2’-endo conformers of 2’dNTPs catalytically inert? It is noteworthy that the sugars of the attacking and substrate nucleotides adopt the 3’-endo conformation in all RNAPs and DNAPs during the nucleotide incorporation ^4^. In other words, even the 3’ ends of DNA primers adopt the 3’-endo conformation to catalyze the incorporation of the 2’dNMPs into the DNA. Apparently, the A-form geometry is much better suited for the catalysis of the nucleotide condensation than the B-form geometry ^3,33^. The better accessibility of the nucleophilic 3’OH group of the attacking nucleotide is likely the primary reason. The substrate then adopts the 3’-endo conformation to match the overall geometry of the A-form duplex and to avoid clashes with the attacking nucleotide ^4^.

In general terms, the inertness of the 2’-endo conformation of the 2’dNTPs can be partially attributed to the differences in the conformations of the triphosphate moieties that in turn originate from the differences in the bond angles at C’4 of the sugar between the 3’- and 2’-endo conformers (**Fig. 1b**). We term this inhibitory component as C’4-geometry-dependent effects. However, in our view, it is impossible to further refine this hypothesis at present: (*i*) the resolutions of the structures are not very high (≥ 2.9 Å, **Supplementary Table 5**) and, (*ii*) the triphosphate moieties are relatively flexible as is evident from the superimposition of different structures (Figure 4 in Ref. ^16^). Moreover, the catalysis requires the isomerization of the triphosphate moiety of the substrate from the pre-insertion into the insertion conformation ^21^, and the structure of the transition state is unknown. High-resolution and time-resolved studies of nucleotide incorporation by the multi-subunit RNAPs would be necessary to determine the reasons behind the inertness of the 2’-endo conformers of 2’dNTPs.

Noteworthy, the conserved Arg is one of only five catalytic residues that are conserved in the superfamily of the so called “two-β-barrel” RNAPs ^34,35^ that includes the multi-subunit RNAPs and very distantly related cellular RNA-dependent RNAPs (RdRps) involved in the RNA interference (**Supplementary Fig. 10**). Accordingly, the common ancestor of the two-β-barrel RNAPs could conceivably discriminate against 2’dNTPs and therefore likely evolved in the presence of both NTPs and 2’dNTPs. This inference lends credence to the hypothesis that proteins evolved in primordial lifeforms that already possessed both RNA and DNA ^36,37^.

Viral RdRps (members of the so-called “right-hand” superfamily of nucleic acid polymerases) are not homologous to multi-subunit RNAPs but share some elements of their sugar selection strategies. It appears that the 3’OH of the substrate NTP facilitates the active site closure in both classes of enzymes. In multi-subunit RNAPs, 3’OH facilitates the TL folding via the interaction with β’Gln929 ^19^, whereas in viral RdRps, 3’OH initiates the closure by sterically clashing with Asp238 (poliovirus RdRp numbering) ^38^. In both classes of enzymes, 2’dNTPs adopt a 2’-endo pose wherein the 3’OH is misplaced and cannot readily facilitate the closure of the active site, explaining low reactivities of 2’dNTPs. However, 3’dNTPs are better substrates than 2’dNTPs also for viral RdRps ^30^ suggesting that the low reactivity of the 2’-endo 2’dNTPs additionally relies on C’4-geometry-dependent effects (see above), which lead to a suboptimal conformation of the triphosphate moiety, a suboptimal geometry of the transition state, or both.

Multi-subunit RNAPs and viral RdRps converged on using the 2’-endo binding pose to discriminate against 2’dNTPs. In doing so these enzymes accentuate the intrinsic preferences of 2’dNTPs to retain the inert 2’-endo conformation upon binding to the A-form template in the non-enzymatic system ^3^. However, the exact implementations of the selection mechanisms are distinct in the two classes of RNAPs. In multi-subunit RNAPs, the 2’-endo pose is stabilized by 3’OH/β’Arg425 attraction, whereas in viral RdRps, the 2’-endo pose is imposed by 3’OH/Asp238 steric clash ^38^. Multi-subunit RNAPs employ the conformational selection and preferentially sample the catalytically-labile 3’-endo conformers of NTP and the catalytically-inert 2’-endo conformers of 2’dNTPs. In contrast, viral RdRps likely rely exclusively on the induced fit ^30^ and bind the catalytically-inert 2’-endo conformers of either NTPs or 2’dNTPs. Following the initial binding, only NTPs can efficiently isomerize into catalytically-labile 3’-endo conformers, ultimately repositioning the Asp238 and switching the RdRp active site on ^38^. These principal differences in the substrate selection mechanisms can be potentially exploited for the concept-based design of sugar-modified substrate analogue inhibitors of viral RdRps such as Remdesivir, one of few drugs currently available for COVID-19 treatment ^39,40^.

In summary, our data show that a universally conserved Arg residue plays a central role in selecting NTPs over 2’dNTPs by the multi-subunit RNAPs. When NTP binds in the RNAP active site, its ribose adopts the 3’-endo conformation that positions the 3’OH group to interact with the universally conserved Gln residue of the TL domain and promotes the closure of the active site, whereas the triphosphate moiety can undergo rapid isomerization into the insertion conformation leading to efficient catalysis. The interaction of the conserved Arg residue with the 2’OH of the NTPs selectively enhances their binding more than 100-fold and renders RNAP saturated with NTPs in the physiological concentration range. In contrast, the interaction of the conserved Arg with the 3’OH of the 2’dNTP substrates shapes their deoxyribose moiety into the catalytically inert 2’-endo conformation where the 3’OH cannot promote closure of the active site and substrate incorporation is additionally inhibited by the unfavorable geometry of the triphosphate moiety. The deformative action of the conserved Arg on the 2’dNTP substrates is an elegant example of active selection against a substrate that is a substructure of the correct substrate.

## Methods

### Reagents and oligonucleotides

DNA and RNA oligonucleotides were purchased from Eurofins Genomics GmbH (Ebersberg, Germany) and IBA Biotech (Göttingen, Germany). DNA oligonucleotides and RNA primers are listed in **Supplementary Table 1**. NTPs, 2’dATP, 3’dGTP and cytidine-5’-[(α,β)-methyleno]triphosphate (CMPCPP) were from Jena Bioscience (Jena, Germany); 2’dGTP, 2’dUTP and 2’dCTP were from Bioline Reagents (London, UK). TECs extended with 3’dGMP did not extend further upon the addition of the next substrate NTP suggesting that 3’dGTP stocks were free of GTP. TECs extended with 2’dGMP, 2’dATP and 2’dUTP migrated faster in the denaturing PAGE than TECs extended with the corresponding NMPs suggesting that 2’dGTP, 2’dATP and 2’dUTP stocks were free of the corresponding NTPs. 2’dCTP stocks were slightly contaminated by CTP as evident from the wild-type and β’M932A RNAPs gels in **Supplementary Figure 7**. These low *K*_*D*_ RNAPs scavenged and depleted the trace amounts of CTP when transcribing the first CMP encoding position but incorporate exclusively 2’dCMP when transcribing CMP encoding positions further downstream. While it was possible to deplete the contaminating CTP by pre-treatment with the unlabeled TEC, we opted to present the experiment with a slightly contaminated 2’dCTP as a showcase of our capabilities to detect contaminations of 2’dNTPs with NTPs.

### Proteins

*E. coli* RNAPs were expressed in the *E. coli* strain T7 Express lysY/Iq (New England Biolabs, Ipswich, MA, USA) and purified by Ni-, heparin and Q-sepharose chromatography as described previously ^41^. RNAPs were dialyzed against storage buffer (50% glycerol, 20 mM Tris-HCl pH 7.9, 150 mM NaCl, 0.1 mM EDTA, 0.1 mM DTT) and stored at −20°C. Plasmids used for protein expression are listed in **Supplementary Table 2**. *T. thermophilus* RNAP holoenzyme was prepared as described previously ^19^.

### TEC assembly

TECs were assembled by a procedure developed by Komissarova et al. ^42^. An RNA primer was annealed to the template DNA, and incubated with 1.5 µM RNAP for 10 min at 25 °C in TB10 buffer (10 mM MgCl2, 40 mM HEPES-KOH pH 7.5, 80 mM KCl, 5% glycerol, 0.1 mM EDTA, and 0.1 mM DTT) and with 2 µM of the non-template DNA for 20 min at 25 °C. For TECs used in nucleotide addition measurements, RNA was the limiting component at 1 µM (final concentration), and the template strand was used at 1.4 µM, whereas for TECs used in the translocation assay the template strand was limiting at 1 µM, and RNA was added at 1.4 µM.

### *In vitro* transcription reactions, processive transcript elongation

The transcription reactions were initiated by the addition of 10 µl of the assembled TEC (0.45 µM) to 10 µl of the substrate mixture (100 µM of each NTP or 2’dNTP), both solutions were prepared in TB10 buffer. In total, five mixtures containing NTPs and 2’dNTPs in different combinations were employed. Four chase mixtures contained three NTPs and one 2’dNTP (2’dATP-, 2’dCTP-, 2d’GTP- and 2’dUTP-chase) whereas the control chase mixture contained four NTPs. The final concentration of NTPs and 2’dNTPs in the reaction mixtures was 50 µM each. The reactions were incubated for 2 min at 25°C and quenched with 40 µl of Gel Loading Buffer (94% formamide, 20 mM Li_4_-EDTA and 0.2% Orange G). RNAs were separated on 16% denaturing polyacrylamide gels and visualized with an Odyssey Infrared Imager (Li-Cor Biosciences, Lincoln, NE, USA); band intensities were quantified using the ImageJ software ^43^.

### Time-resolved nucleotide addition measurements

Time-resolved measurements of nucleotide addition were performed in an RQF 3 quench-flow instrument (KinTek Corporation, Austin, TX). The reaction was initiated by the rapid mixing of 14 µl of 0.2 µM TEC with 14 µl of GTP (400, 2000 or 4000 µM) or 2’dGTP (400, 2000 or 4000 µM) or 3’dGTP (5, 20, 50, 200, 1000 µM). Both TEC and substrate solutions were prepared in TB10 buffer. The reaction was allowed to proceed for 0.004–300 s at 25°C and quenched by the addition of 86 µl of 0.45 M EDTA or 0.5 M HCl. EDTA quenched reactions were mixed with 171 µl of Gel oading Buffer. HCl quenched reactions were immediately neutralized by adding 171 µl of Neutralizing-Loading Buffer (94% formamide, 290 mM Tris base, 13 mM Li4-EDTA, 0.2% Orange G). RNAs were separated on 16% denaturing polyacrylamide gels and visualized with an Odyssey Infrared Imager (Li-Cor Biosciences, Lincoln, NE, USA); band intensities were quantified using the ImageJ software ^43^.

### Time-resolved fluorescence measurements

Measurements were performed in an Applied Photophysics (Leatherhead, UK) SX.18MV stopped-flow instrument at 25°C. The 6-MI fluorophore was excited at 340 nm and the emitted light was collected through a 400 nm longpass filter. The nucleotide addition reactions were initiated by mixing 60 µl of 0.2 µM TEC in TB10 buffer with 60 µl of GTP (5 - 4000 µM) or 2’dGTP (100 - 4000 µM) in TB10 buffer. At least three individual traces were averaged for each concentration of the substrate.

### Data analyses

Time-resolved GMP incorporation data (HCl and EDTA quenched reactions) and the translocation timetraces were simultaneously fitted to a three-step model using the numerical integration capabilities of the KinTek Explorer software ^44^ (KinTek Corporation, Austin, TX) largely as described previously ^45^. The model postulated that the initial TEC16 reversibly binds the GTP substrate, undergoes the irreversible transition to TEC17 upon incorporation of the nucleotide into RNA, followed by the irreversible translocation. The EDTA quenched reactions were modeled using the pulse-chase routine of the Kin-Tek Explorer software. Time-resolved 2’dGMP incorporation concentration series (translocation timetraces) were globally fitted to a stretched exponential function (**Equation 1, Supplementary Table 4**): the exponent followed a hyperbolic dependence on the 2’dGTP concentration; *Km*, rate constant *k* and the stretching parameter β were shared by all curves in the dataset. A detailed description of the data analyses is presented in **Supplementary Note**.

### Docking experiments

RNAP fragments comprising amino acid residues and template DNA within 20 Å from the active-site-bound substrates were extracted from the X-ray crystal structures of the initiation (PDB ID 4Q4Z) ^19^ and initially transcribing (PDB ID 6WOX) (this work) complexes of *T. thermophilus* RNAP. The substrate binding sites were vacated by removing CMPCPP (RNAP fragment 1) and 2’dCTP (RNAP fragment 2). The three-dimensional structures of the 3’-endo CTP, 3’-endo 2’dCMPNPP and 2’-endo 2’dCMP were extracted from PDB ID 3BSO (1.74 Å) ^46^, 4O3N (1.58 Å) ^47^ and 3FL6 (1.17 Å) ^48^, respectively. Phosphate moieties of the ligands were rebuilt using Discovery Studio 4.5 (Accelrys, San Diego, CA, USA) to produce 3’-endo CTP, CMP, 2’dCTP, 2’dCMP, 2’-endo 2’dCTP and 2’dCMP. Ligands and RNAP fragments were prepared for docking using AutoDock tools ^49^. AutoDock Vina 1.1.2 docking runs were performed in a 16 × 20 × 20 Å search space centered at 183, 6, 83 Å (coordinate space of the RNAP fragment 1) or in a 30 × 30 × 30 Å search space centered at 460, 9, 670 Å (coordinate space of the RNAP fragment 2) using the default scoring function ^50^. The docking was performed using 12 simultaneous computational threads; 20 binding poses were recorded for each run. Binding poses involving Watson-Crick pairing between the substrate and the template DNA (templated poses) were manually selected and extracted for further analysis. The position of the sugar moiety in templated poses closely matched that observed in crystal structures because any significant rotation around the glycosidic bond, the only available degree of freedom (for templated poses), would lead to steric clashes.

Our initial docking trials revealed that docking of nucleoside monophosphates produced the most robust and quantitatively interpretable results. Thus, the docking algorithm failed to recover templated poses for nucleosides without phosphate groups. The docking algorithm also failed to position the triphosphate moiety to coordinate metal ion number two and instead attempted to maximize its contacts with the protein. As a result, the recovered conformations of the triphosphate moieties differed from those observed in crystal structures. Considering the high impact of the triphosphate moiety on the ligand binding score and our assessment that the triphosphate moiety was docked incorrectly, we opted to limit the systematic investigation of the interaction between RNAP and the sugar moieties of nucleosides to docking nucleoside monophosphates.

We first docked 3’-endo CMP, 3’-endo 2’dCMP and 2’-endo 2’dCMP to the RNAP fragment 1. The docking algorithm recovered high-scoring poses (−7.8 ±0.1 kcal/mol) for CMP in 10 out of 10 runs, lower-scoring poses (−6.8 ±0.2 kcal/mol) for 3’-endo 2’dCMP in 8 out of 10 runs and 2’-endo 2’dCMP in 5 out of 10 runs. The β’Arg425 side chain was kept flexible in the latter case because our manual assessment suggested that a sub-angstrom repositioning of β’Arg425 would be needed to accommodate the 2’-endo deoxyribose. We than fixed the β’Arg425 conformation observed in one of the highest scoring poses and performed 10 additional docking runs. This time templated poses (−7.0 ±0.1 kcal/mol) were recovered in 10 out of 10 runs (**Supplementary Table 6**). These *in silico* experiments suggested that the semi-closed active site can bind the 3’-endo and 2’-endo 2’dCMP with similar affinities. The 3’OH of the 3’-endo 2’dCMP was positioned to interact with β’Gln929 and β’Asn458, whereas the 3’OH of the 2’-endo 2’dCMP was positioned to interact with β’Arg425 and β’Asn458 (**Supplementary Fig. 8**). We further inferred that the open active should have preference for the 2’-endo 2’dCMP because β’Gln929 is not positioned to interact with the 3’OH of the substrate in the open active site. Well in line with our prediction, the X-ray diffraction data for the crystals of the RNAP-2’dCTP complex was consistent with the 2’-endo conformation of the 2’dCTP bound in the open active site (**Fig. 4b, c, Supplementary Fig. 9a**). We further verified the binding preferences of the open active site *in silico* by removing the 2’dCTP from the model and docking alternative conformers of the 2’dCMP. The docking algorithm recovered higher-scoring poses (−6.6 ±0.1 kcal/mol) for the 2’-endo 2’dCMP in 9 out of 10 runs and lower-scoring poses (−6.3 ±0.1 kcal/mol) for 3’-endo 2’dCMP in only 2 out of 10 runs (**Supplementary Table 6**).

### Preparation of the promoter DNA scaffold for the crystallization

The non-template DNA strand (5’-TATAATGGGAGCTGTCACGGATGCAGG-3’) was annealed to the template DNA strand (5’-CCTGCATCCGTGAGTGCAGCCA-3’) in 40 µl of 10 mM Tris-HCl (pH 8.0), 50 mM NaCl, and 1 mM EDTA to the final concentration of 1 mM. The solution was heated at 95 °C for 10 min and then gradually cooled to 22 °C.

### Crystallization of the *T. thermophilus* RNAP initially transcribing complexes

The crystals of the RNAP and promoter DNA complex were prepared as described previously ^51,52^. The RNAP and promoter DNA complex was prepared by mixing 24 µl of 18 µM *T. thermophilus* holoenzyme (in 20 mM Tris-HCl, pH 7.7, 100 mM NaCl, and 1% glycerol) and 0.65 µl of 1 mM DNA scaffold and incubated for 30 min at 22 °C. Crystals were obtained by using hanging drop vapor diffusion by mixing equal volume of RNAP-DNA complex solution and crystallization solution (100 mM Tris-HCl, pH 8.7, 200 mM KCl, 50 mM MgCl_2_, 10 mM Spermine tetra-HCl, and 10% PEG 4000) and incubating at 22 °C over the same crystallization solution. The crystals were cryoprotected by soaking in same constituents as the crystallization solution with stepwise increments of PEG4000 and (2R,3R)-(—)-2,3-butanediol (Sigma-Aldrich) to final concentrations of 25 and 15%, respectively. The crystals were sequentially transferred to the final cryoprotection solution containing 1 mM primer 5’-GpCpA-3’ and either 4 mM 2’dCTP or 3’dCTP. The crystals were harvested and flash frozen in liquid nitrogen.

### X-ray data collections and structure determinations

The X-ray datasets were collected at the Macromolecular Diffraction at the Cornell High Energy Synchrotron Source (MacCHESS) F1 beamline (Cornell University, Ithaca, NY) and structures were determined as previously described ^51,52^ using the following crystallographic software: HKL2000 ^53^, Phenix ^54^ and Coot ^55^.

## Data availability

Atomic coordinates and structure factors for the reported crystal structures have been deposited with the Protein Data bank under accession number 6WOX and 6WOY.

## Contributions

J.J.M. and E.V. performed biochemical analysis of *E. coli* RNAPs. K.S.M and Y.S. solved X-ray structures of *T. thermophilus* RNAP. G.A.B performed *in silico* docking experiments. All authors contributed to the analysis of the data and the interpretation of the results. G.B. and K.S.M wrote the manuscript with contributions from the other authors.

## Acknowledgement

We thank Irina Artsimovitch for critically reading the manuscript, the staff at the MacCHESS for support of crystallographic data collection, Anssi M. Malinen for constructing plasmids, Matti Turtola for his contribution to the development of the EDTA quench method.

---

**Supplementary Fig. 1:**
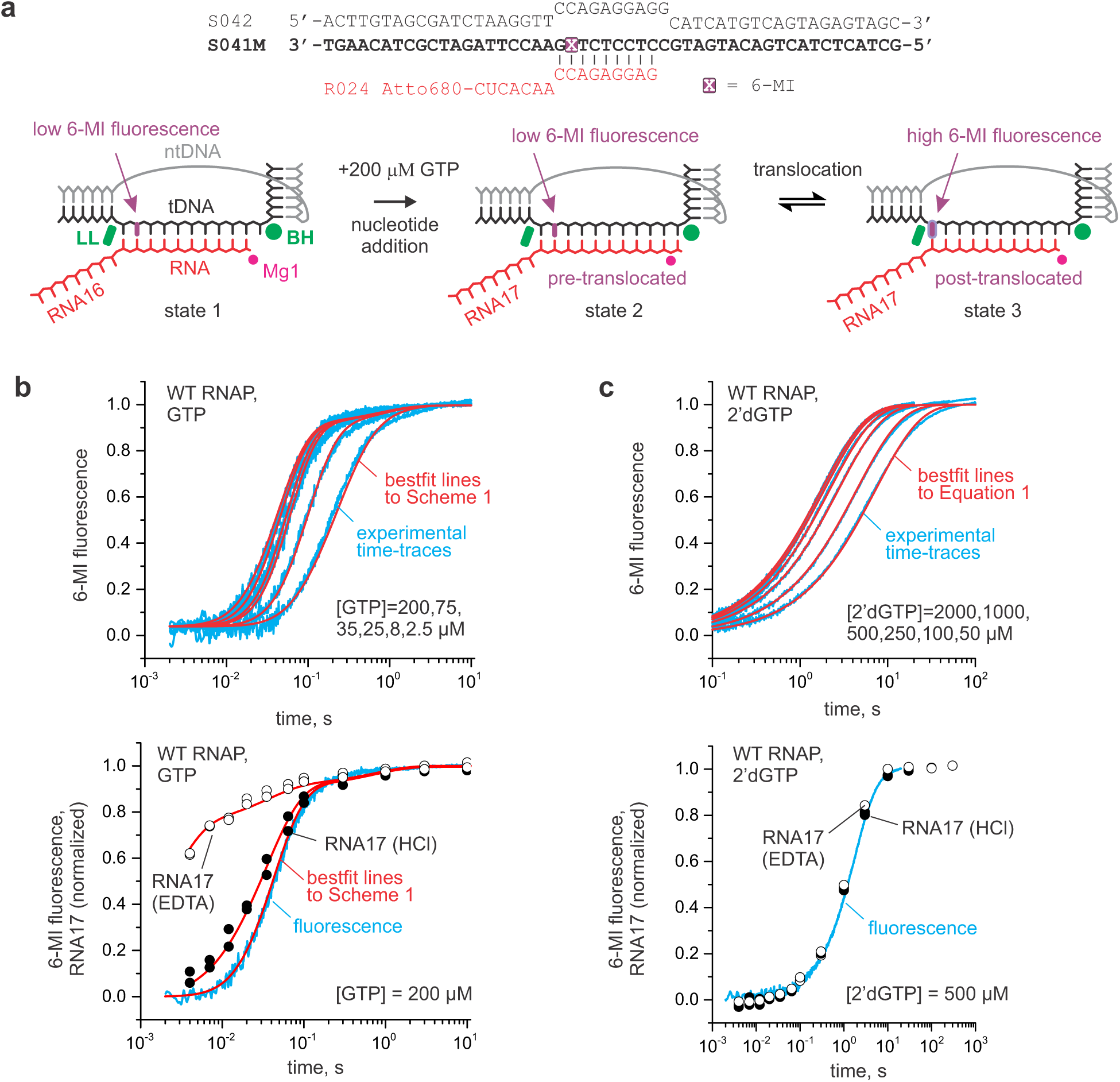
Concentration series of GMP and 2’dGMP incorporation by the WT *E. coli* RNAP. **a** The nucleic acid scaffold employed in translocation and nucleotide addition assays. The fluorescence of a guanine analogue 6-MI was quenched by the neighboring base pairs in the initial TEC (*state 1*) and the pre-translocated TEC that formed following the nucleotide incorporation (*state 2*) but increased when the 6-MI relocated to the edge of the RNA:DNA hybrid upon translocation (*state 3*). The β’ Bridge Helix (BH) and the β’ Lid Loop (LL) flank the RNA:DNA hybrid in the multi-subunit RNAPs. **b** GMP incorporation. *Top panels*: Concentration series. *Bottom panels*: EDTA or HCl quenched reactions and a fluorescent time-trace at 200 µM GTP. The data in were fit to **Scheme 1. c** 2’dGMP incorporation. *Top panels*: Concentration series. *Bottom panels*: EDTA or HCl quenched reactions and a fluorescent time-trace at 500 µM 2’dGTP. The data were fit to **Equation 1**.

**Supplementary Fig. 2:**
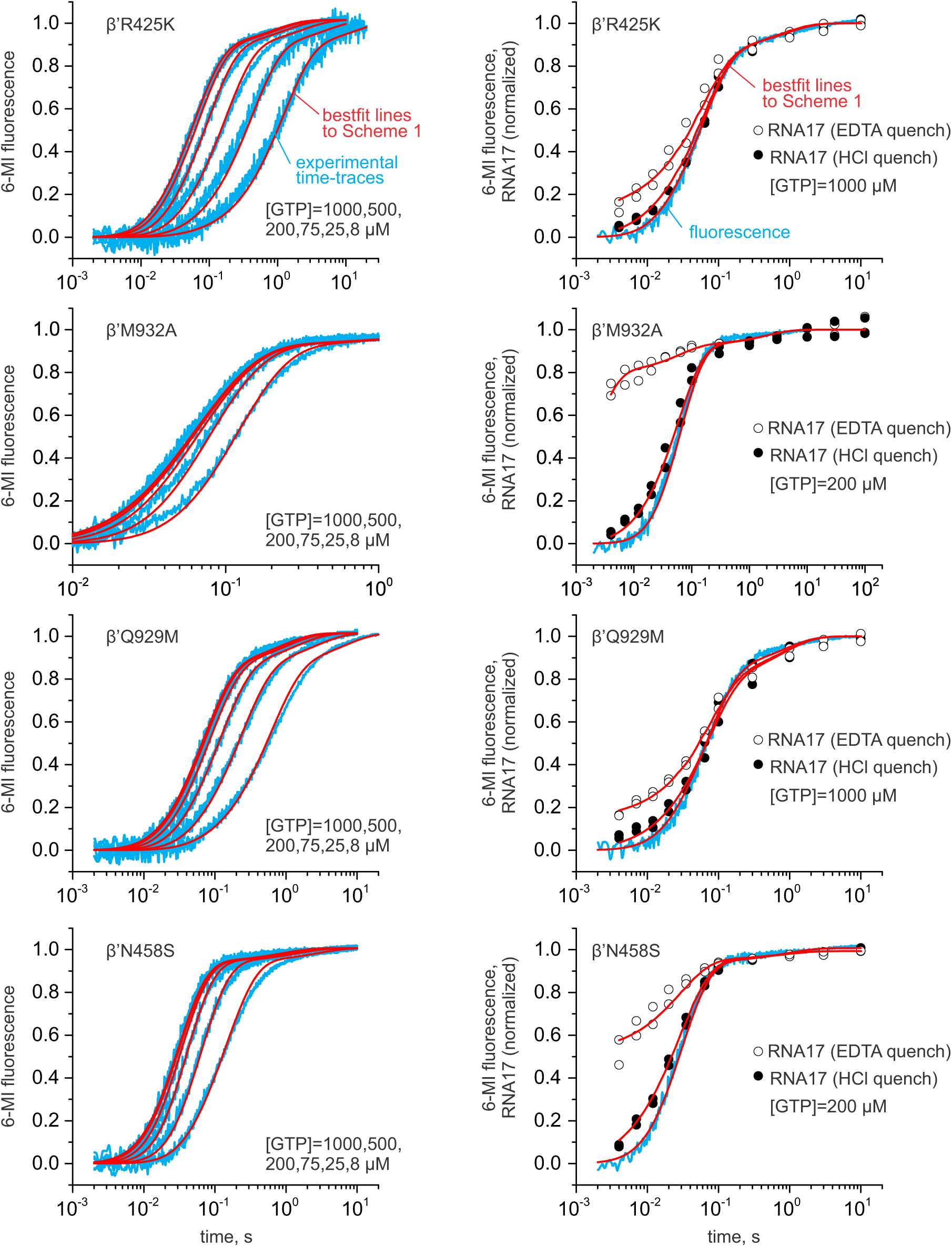
Concentration series of GMP incorporation by the variant *E. coli* RNAPs. *Left graphs:* fluorescence time-traces (blue) at six GTP concentrations. *Right graphs:* EDTA or HCl quenched reactions and a fluorescent timetrace at 200 µM (β’M932A and β’N458S) or 1000 µM (β’R425K and β’Q929M) GTP. Bestfit lines to **Scheme 1** are shown in red.

**Supplementary Fig. 3:**
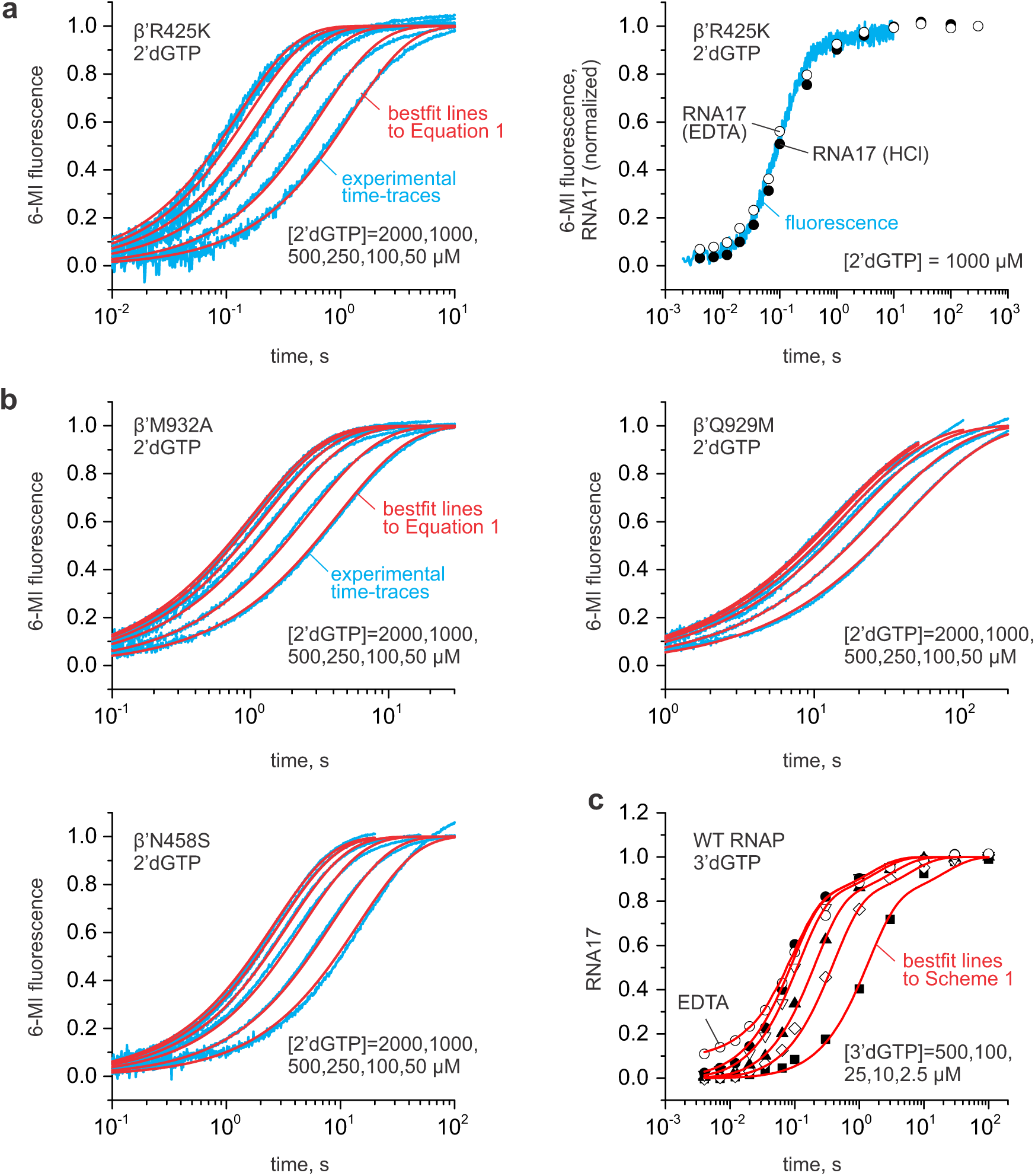
2’dGTP and 3’dGTP concentration series by the WT and variant *E. coli* RNAPs. **a** Incorporation of 2’dGMP by β’R425K RNAP. *Left panel:* Concentration series. *Right panel:* EDTA or HCl quenched reactions and a fluorescent timetrace at 1000 µM 2’dGTP. Best fit lines to **Equation 1** are shown in red. **b** Incorporation of 2’dGMP by β’M932A, β’Q929M and β’N458S RNAPs. **c** Incorporation of 3’dGMP by the wild-type RNAP. RNA extension time curves were obtained by HCl quench at five 3’dGTP concentration (closed circles, triangles, diamonds and squares). RNA extension time curve at 500 µM 3’dGTP was additionally measured using EDTA quench (open circles). Best fit lines to **Scheme 1** are shown in red.

**Supplementary Fig. 4:**
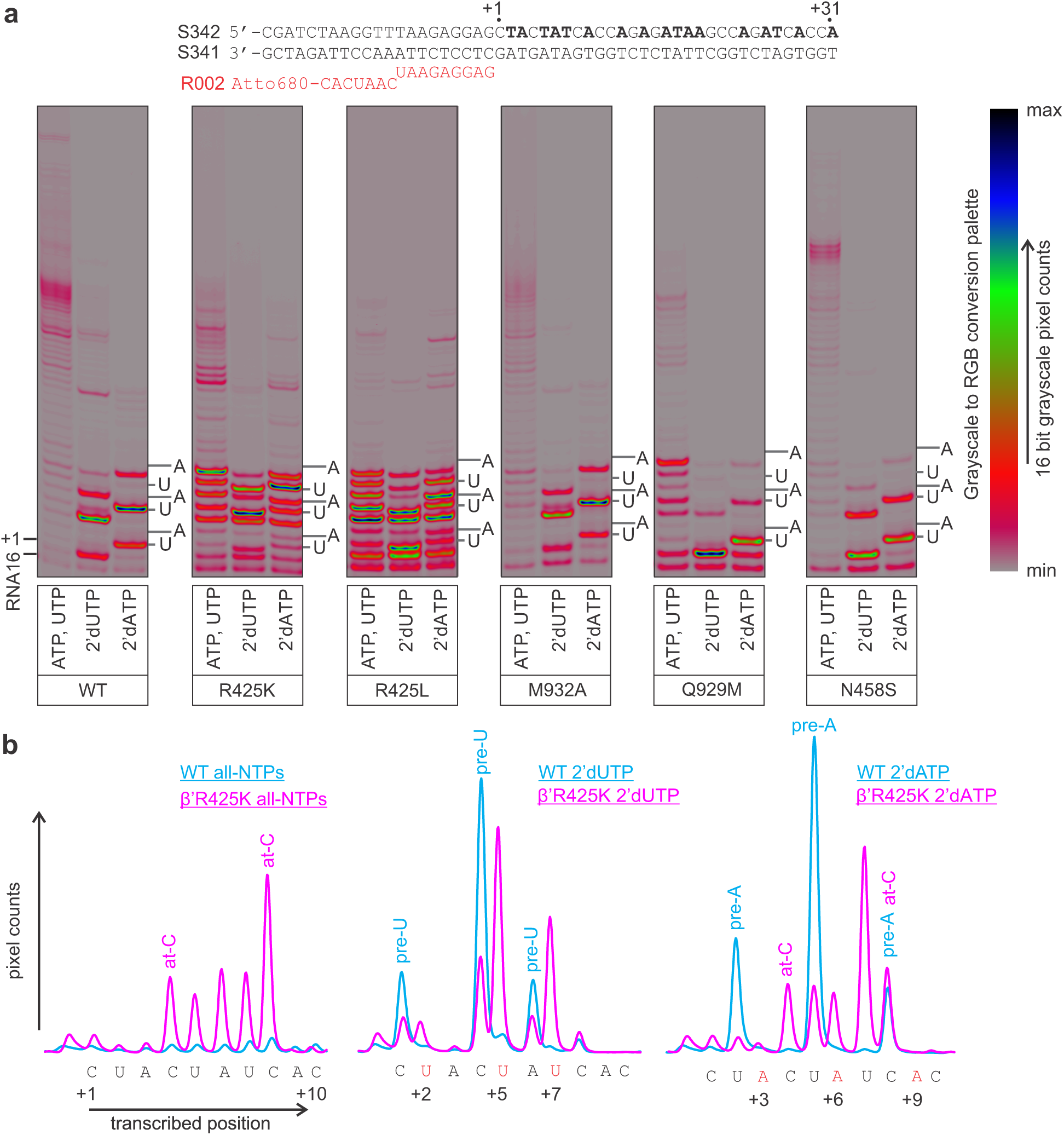
Utilization of 2’dUTP and 2’dATP during the processive transcript elongation by the WT and variant RNAPs. **a** TECs were assembled using the scaffold shown above the gel panels and chased with 100 µM CTP, GTP, UTP, ATP (all-NTPs-chase), or CTP, GTP, ATP, 2’dUTP (2’dUTP-chase), or CTP, GTP, UTP, 2’dATP (2’dATP-chase) for 5 min at 25°C. The positions of UMPs or AMPs in resolved stretches of the transcribed sequence are marked along the right edge of the gel panels. 16-bit grayscale scans were normalized using max pixel counts within each gel panel and pseudocolored using RGB palette on the right. **b** Lane profiles of transcription by the WT (cyan) and β’R425K (magenta) RNAPs quantified from gels in **(a)**. Traces were manually aligned along the X-axis and scaled along the Y-axis using several sequence positions as references.

**Supplementary Fig. 5:**
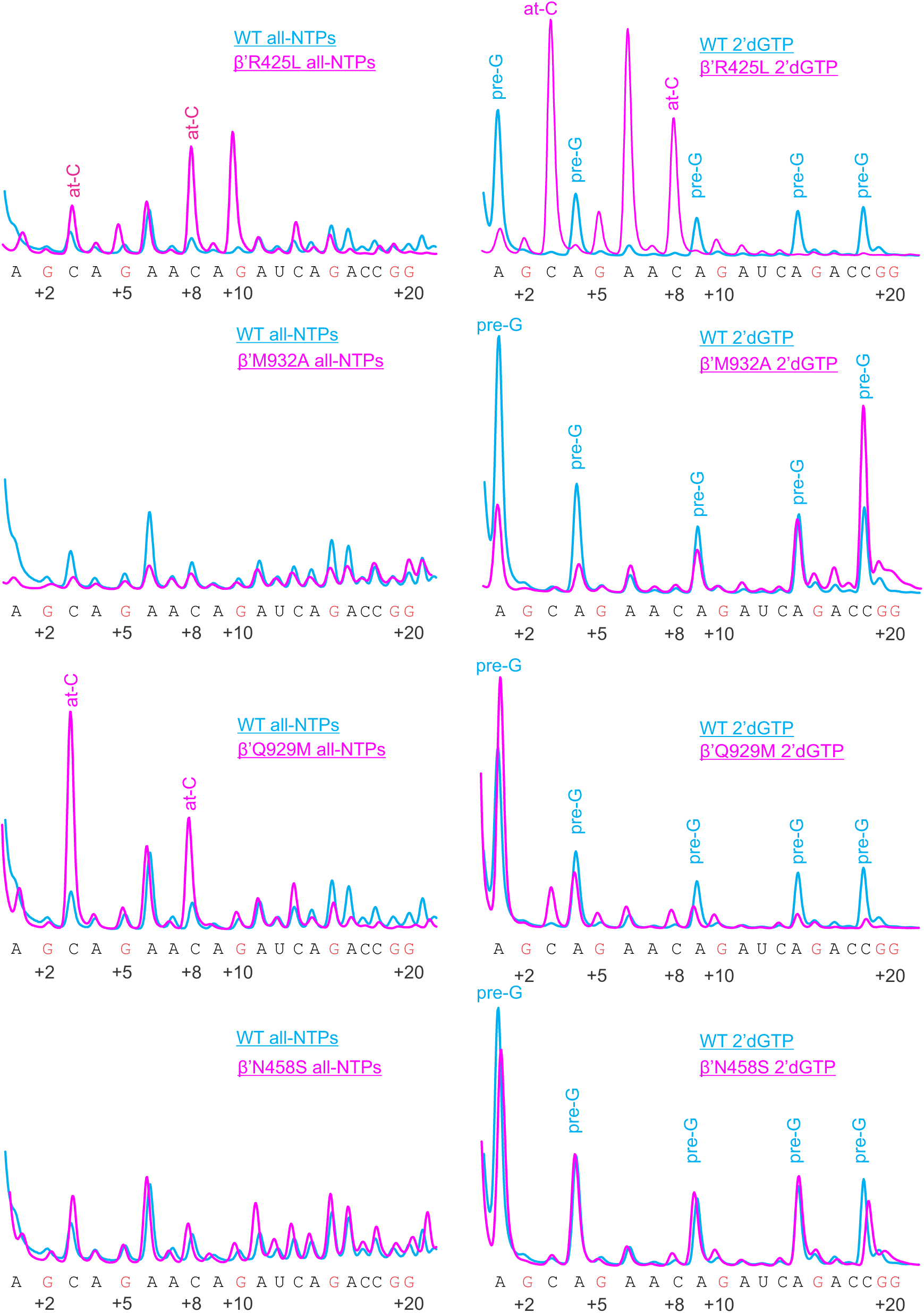
Lane profiles of transcription in all-NTPs and 2’dGTP chases quantified from gels in main text Figure 3.

**Supplementary Fig. 6:**
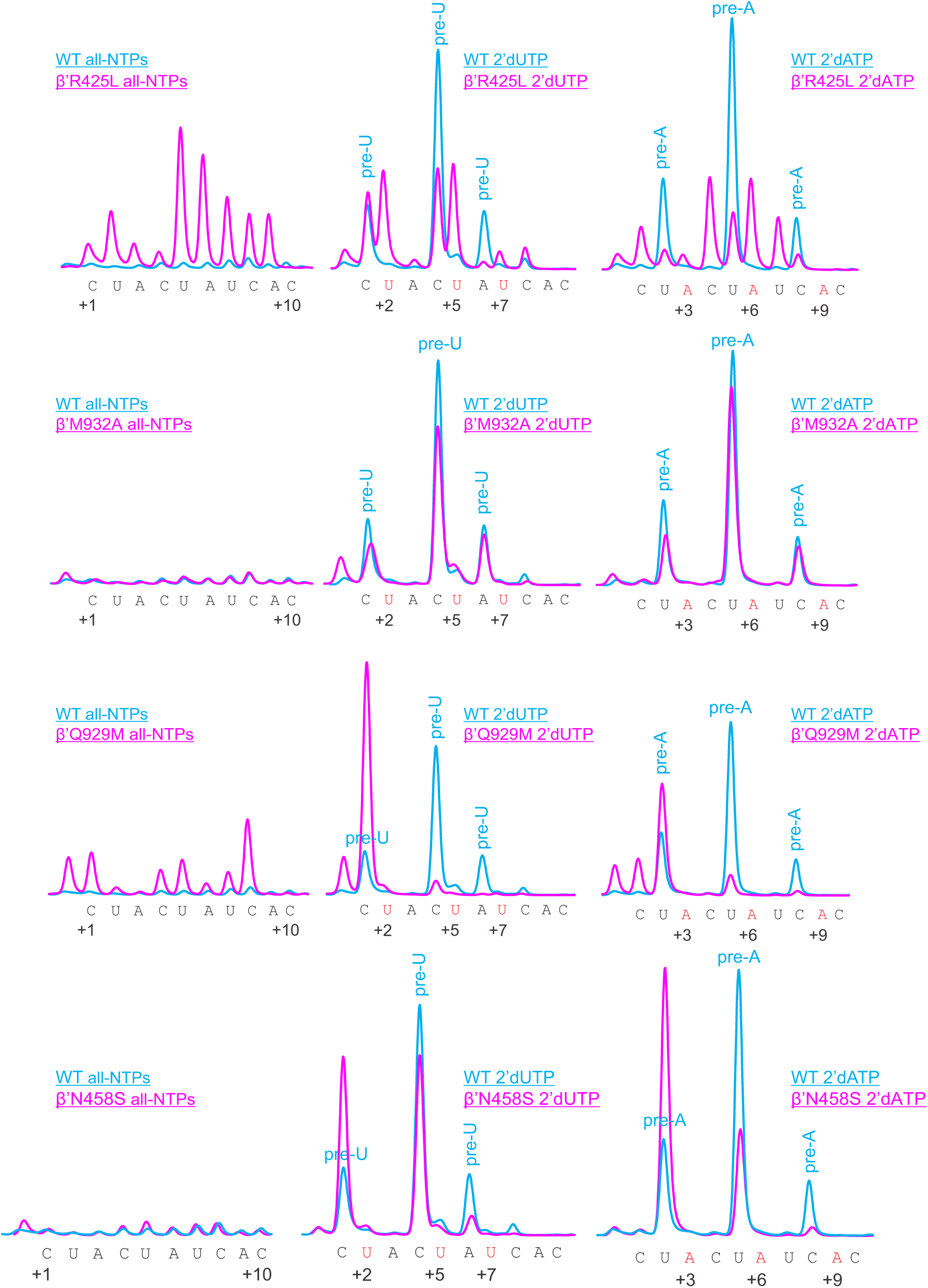
Lane profiles of transcription in all-NTPs, 2’dUTP and 2’dATP chases quantified from gels shown in Supplementary Figure 4.

**Supplementary Fig. 7.**
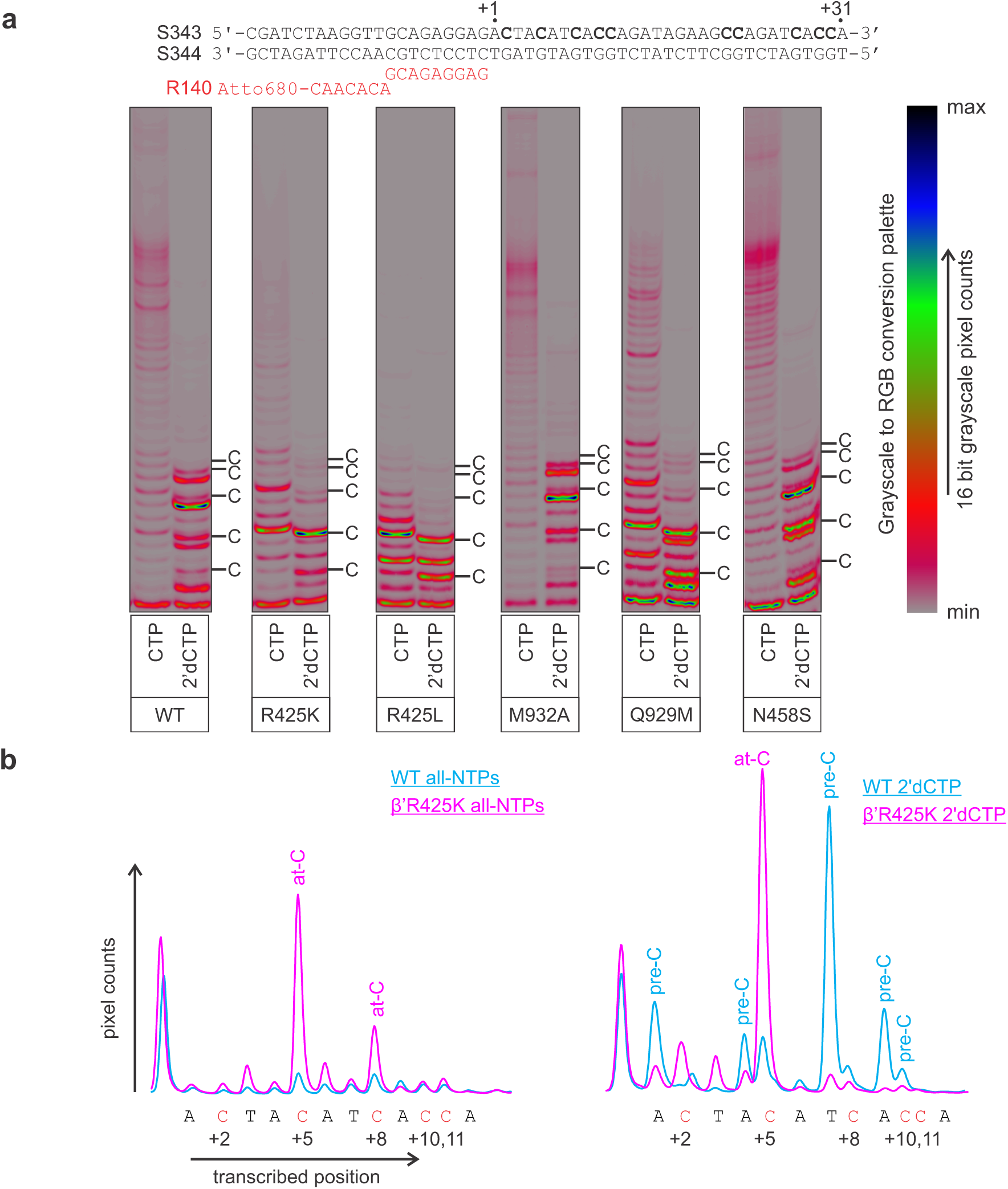
Utilization of 2’dCTP during the processive transcript elongation by the WT and variant RNAPs. **a** TECs were assembled using the scaffold shown above gel panels and chased with 100 µM GTP, UTP, ATP and CTP (all-NTPs chase) or 2’dCTP (2’dCTP chase) for 2 min at 25°C. **b** Lane profiles of transcription by the WT (cyan) and β’R425K (magenta) RNAPs quantified from gels in **(a)**. RNAPs with low *K*_*D*_^*CTP*^ scavenged the trace amounts of CTP when transcribing the first CMP encoding position in 2’dCTP-chase: the WT RNAP produced almost exclusively a slow whereas β’M932A produced a double band. However, RNAPs depleted the contaminating CTP when transcribing the first CMP encoding position and incorporated exclusively 2’dCMP when transcribing CMP encoding positions further downstream.

**Supplementary Fig. 8:**
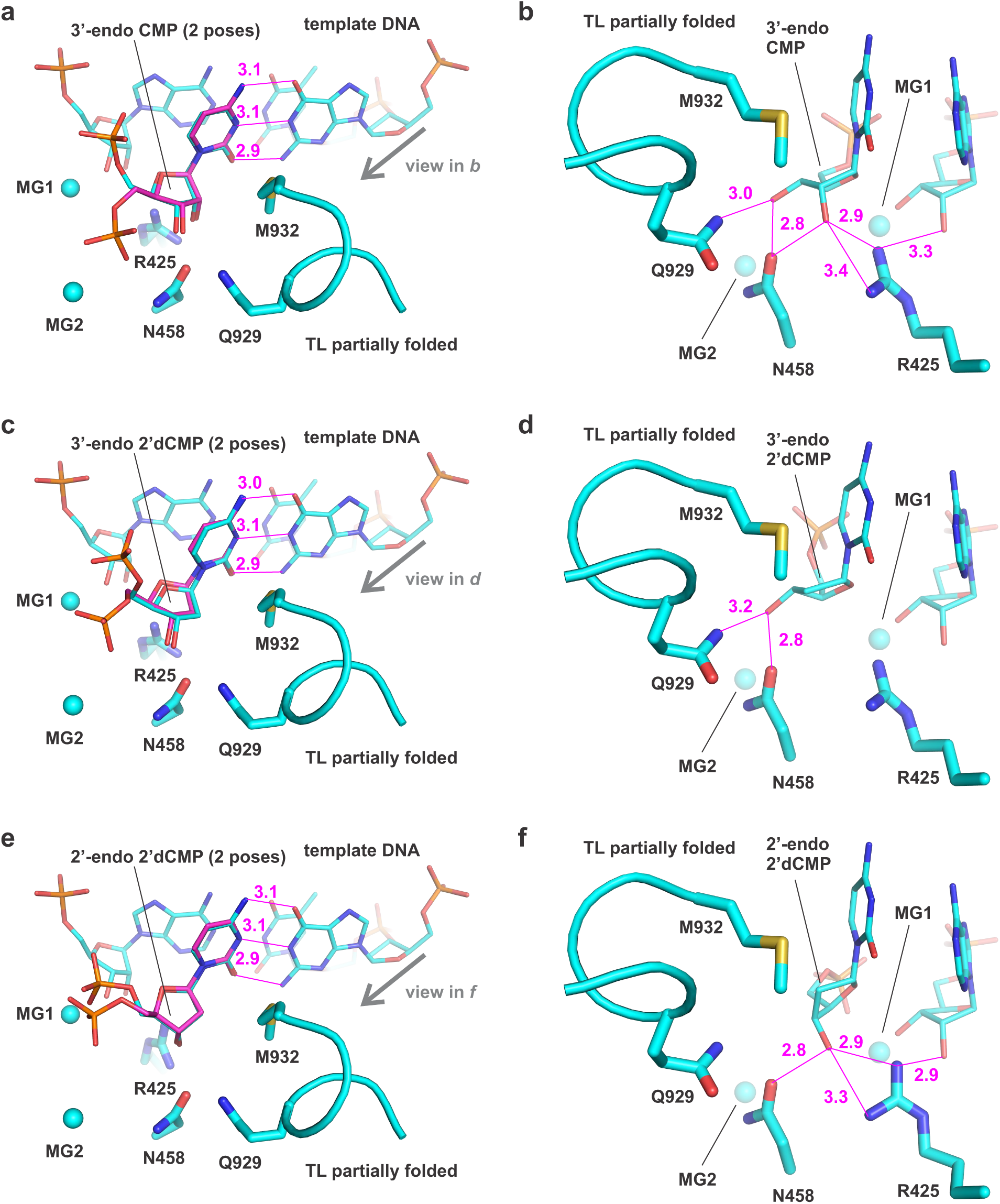
*In silico* docking of 3’-endo CMP (a, b), 3’-endo 2’dCMP (c, d) and 2’-endo 2’dCMP (e, f) in the active site of *T. thermophilus* RNAP. Left panels show two representative poses for each ligand colored cyan and magenta. Right panels show only cyan pose for each ligand. Magenta numbers are interatomic distances in Å.

**Supplementary Fig. 9:**
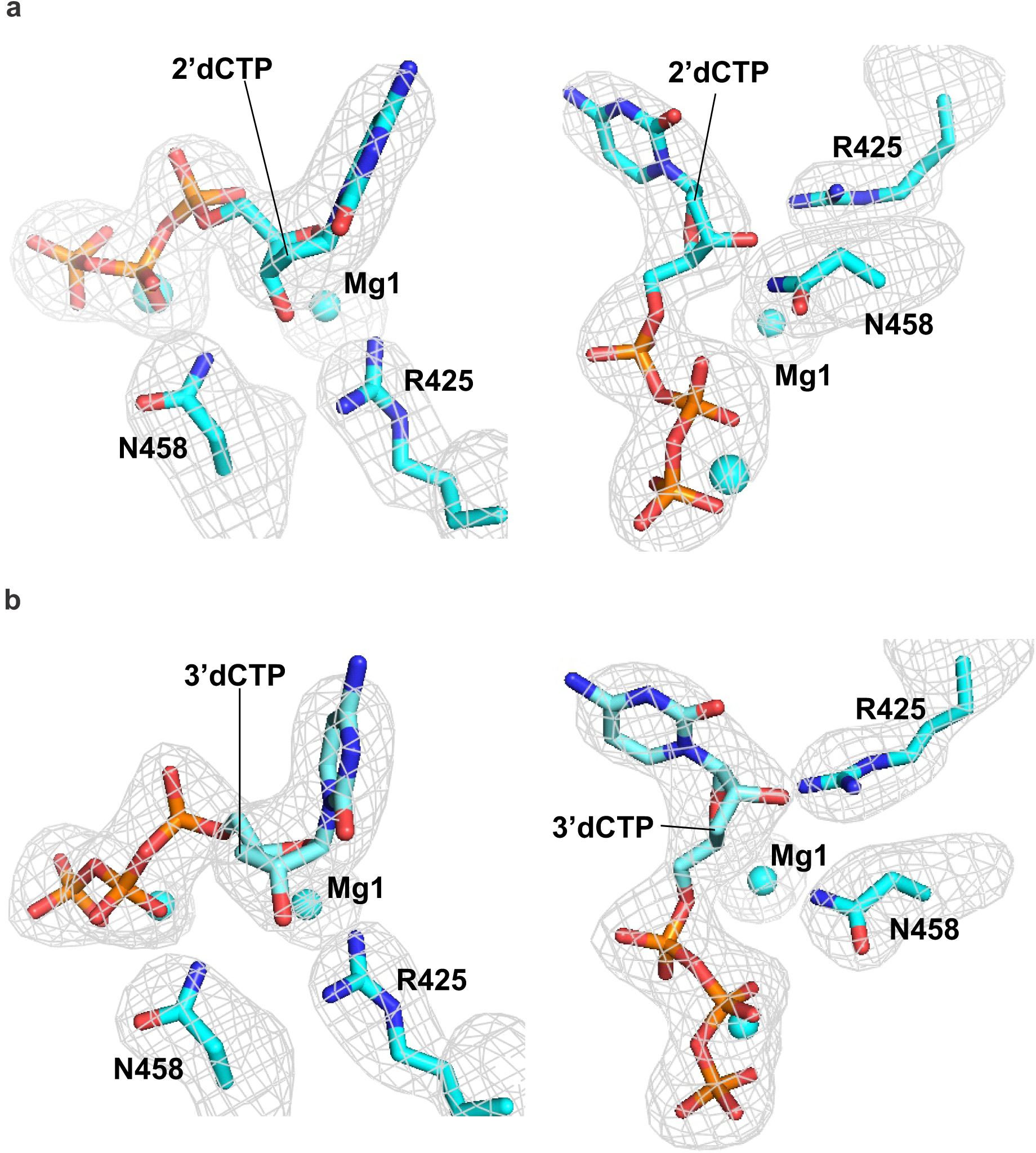
Orthogonal views of 2Fo-Fc electron density maps (gray mesh, 1.0 σ) for the active site regions of *T. thermophilus* RNAP with bound 2’dCTP (a) and 3’dCTP (b). Backgrounds residues were removed for the clarity. 2’dCTP, 3’dCTP and amino acid residues β’R425 and β’N458 are shown as sticks. The Mg^2+^ and Na^+^ ions are shown as cyan spheres.

**Supplementary Fig. 10.**
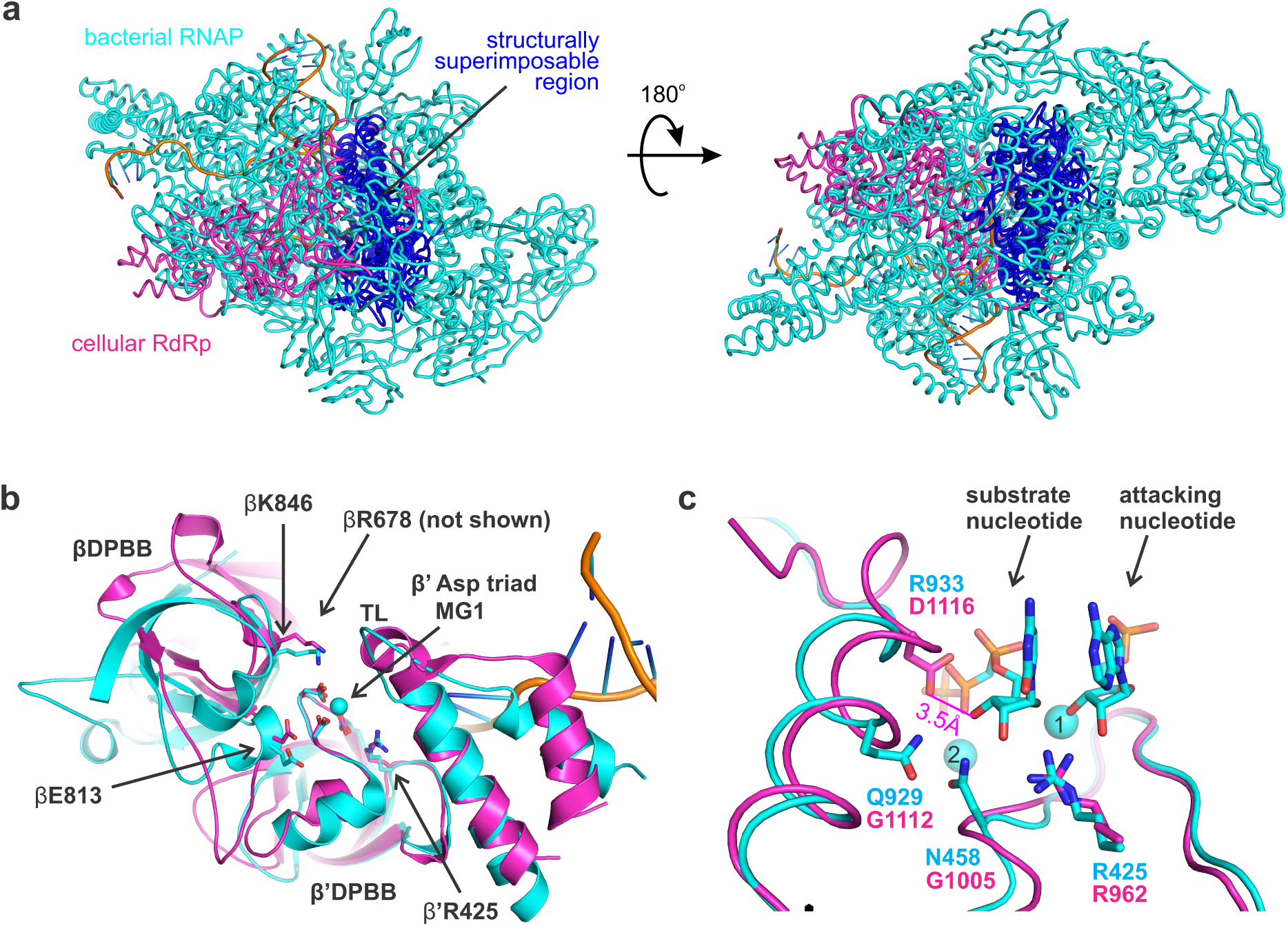
Conservation of the nucleo-sugar recognition in two-β-barrel RNAPs. **a** A superimposition of a multi-subunit RNAP (cyan, PDB ID 4Q4Z) and a monomer of the homodimeric cellular RdRp (magenta, PDB ID 2J7N). Homologous regions are colored blue in both RNAPs. Structures were superimposed using the β’Arg425 and three Asp triad residues as references. **b** A close-up view of the active site. Six out of seven conserved catalytic residues (shown as sticks) are contributed by loops of the double-psi β-barrels (DPBB). In multi-subunit RNAPs, βDPBB and β’DPBB are contributed by β and β’ subunits, respectively, whereas in RdRps both DPBBs belong to the same polypeptide chain. **c** A close-up view centered on the ribose moiety of the substrate. Catalytic Mg^2+^ ions are numbered. The β’Arg425 adopts similar conformations in both RNAPs. In contrast, the β’Asn458 and β’Gln929 correspond to Gly residues in cellular RdRps. Asp1116 of RdRp (corresponds to β’Arg933 in *E. coli* RNAP and Asn1082 in *S. cerevisiae* RNAPII) resides 3.5 Å from the 3’OH of the substrate from the superimposed structure. Asp1116 may therefore play role in the recognition of the 3’OH of substrate NTPs by RdRps.

## Supplementary Data

Interactive 3D versions of the structural figures (WebGL in browser):

**Supplementary Data 1: Interactive Supplementary Fig. 8a,b**

https://belogurov.org/2disc/docked_3endo_CMP.html

**Supplementary Data 2: Interactive Supplementary Fig. 8c, d**

https://belogurov.org/2disc/docked_3endo_2dCMP.html

**Supplementary Data 3: Interactive Supplementary Fig. 8e, f**

https://belogurov.org/2disc/docked_2endo_2dCMP.html

**Supplementary Data 4: Interactive Fig. 4a, b**

https://belogurov.org/2disc/x-ray_cmpcpp.html

**Supplementary Data 5: Interactive Fig. 4c, d**

https://belogurov.org/2disc/x-ray_2dCTP.html

**Supplementary Data 6: Interactive Fig. 4e, f**

https://belogurov.org/2disc/x-ray_3dCTP.html

## Supplementary Tables

**Supplementary Table 1.**
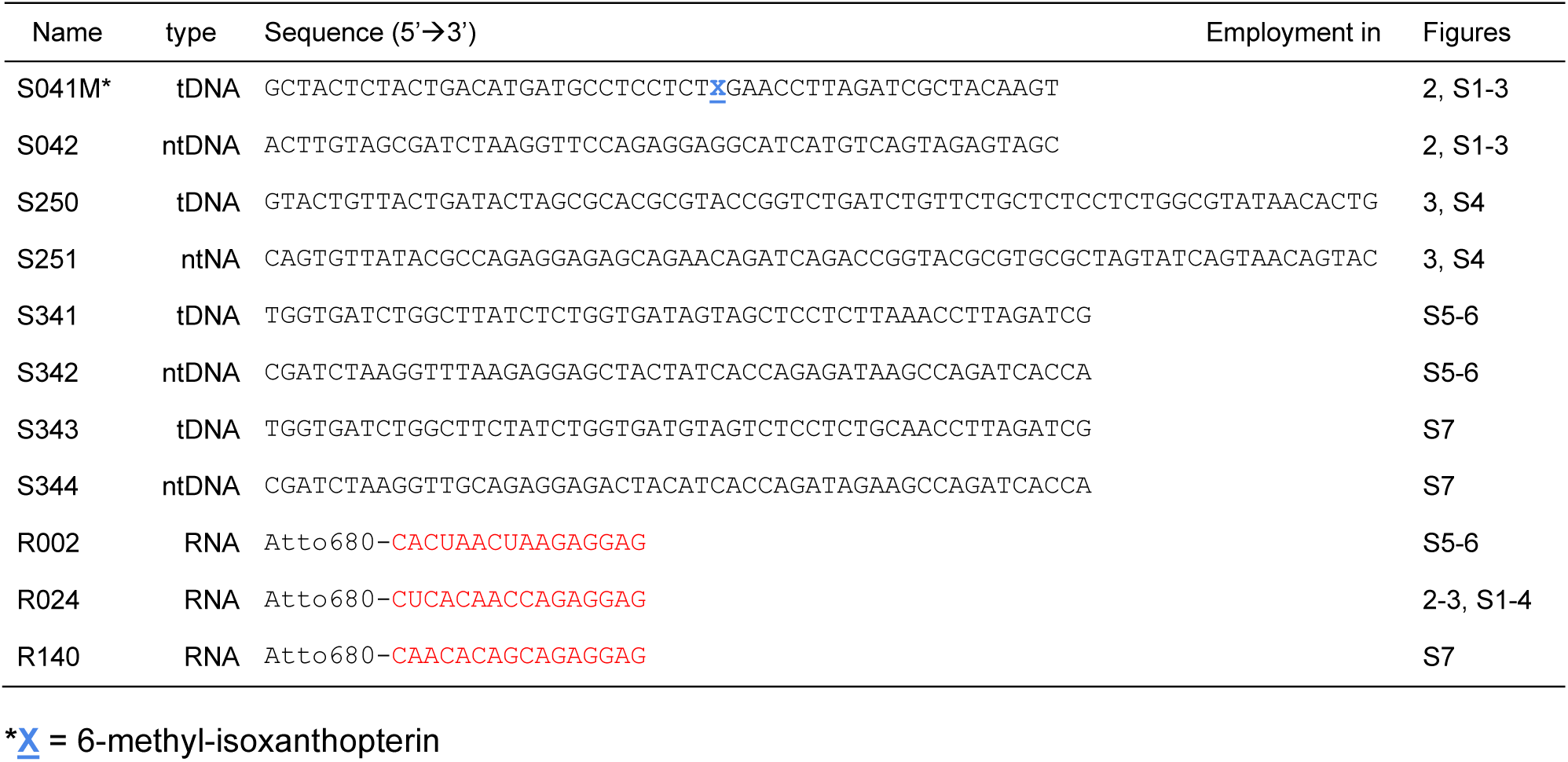
DNA oligonucleotides and RNA primers used in this study.

**Supplementary Table 2.** *E. coli* protein expression vectors used in this study.

| Name | Description | Source/reference |
| --- | --- | --- |
| pVS10 | wild-type RNAP (T7p-α-β-β'-His<sub>6</sub>-ω) | Ref. <sup>41</sup> |
| pAM012 | β'M932A RNAP (T7p-α-β-β'[M932A]_TEV_His<sub>10</sub>-ω) | Ref. <sup>29</sup> |
| pAM017 | β'R425L RNAP (T7p-α-β-β'[R425L]_TEV_His<sub>10</sub>-ω) | this work |
| pAM018 | β'R425K RNAP (T7p-α-β-β'[R425K]_TEV_His<sub>10</sub>-ω) | this work |
| pJM017 | β'Q929M RNAP (T7p-α-β-β'[Q929M]_TEV_His<sub>10</sub>-ω) | this work |
| pIA528+pIA839 | β'N458S RNAP (T7p-α-β-β'[N458S]_His<sub>6</sub>) + *araB*p-ω | Ref. <sup>26</sup> |

**Supplementary Table 3.** Kinetic parameters for the reversible binding of GTP and incorporation of GMP by the wild-type and altered *E. coli* RNAPs.

| RNAP | $k_{\text{on}}$, $\mu\text{M}^{-1}\text{s}^{-1}$ | $k_{\text{off}}$, $\text{s}^{-1}$ | $k_{\text{cat}}$, $\text{s}^{-1}$ (fast) | $k_{\text{cat}}$, $\text{s}^{-1}$ (slow) | fast fraction, % | $k_{\text{tra}}$, $\text{s}^{-1}$ | $K_D$, $\mu\text{M}$ |
| --- | --- | --- | --- | --- | --- | --- | --- |
| WT | 2.5 (2.2 – 3.0) | 7.7 (4.0 – 9.6) | 31 (28 – 42) | 2.6 (0.8 – 7.6) | 88 (77 – 93) | 96 (61 – 140) | 3.1 (2.3 – 4.0) |
| $\beta'$R425K | 0.5 (0.4 – 0.8) | 72 (57 – 120) | 20 (18 – 25) | 1.3 (0.4 – 4.1) | 86 (76 – 92) | 220 (110 – 1600) | 140 (110 – 160) |
| $\beta'$M932A | 2.8 (2.2 – 8.4) | 3.1 (0.8 – 5.4) | 19 (17 – 29) | 3.6 (0.2 – 11) | 86 (45 – 98) | 96 (61 – 230) | 1.1 (0.6 – 1.4) |
| $\beta'$Q929M | 1.1 (0.9 – 1.6) | 62 (49 – 92) | 16 (14 – 20) | 1.3 (0.6 – 2.3) | 79 (73 – 87) | 140 (90 – 430) | 56 (41 – 57) |
| $\beta'$N458S | 1.7 (1.5 – 2.0) | 28 (22 – 35) | 37 (34 – 41) | 1.2 (0.1 – 5.5) | 94 (89 – 96) | 170 (121– 260) | 16 (12 – 18) |

The reaction products were modelled as sums of independent contributions by the fast and slow fractions of RNAP using numerical integration capabilities of the KinTek Explorer software. Contributions of each fraction were modeled as **Scheme 1**. Upper and lower bounds of the parameters were calculated at a 10% increase in $\text{Chi}^2$ .

**Supplementary Table 4.**
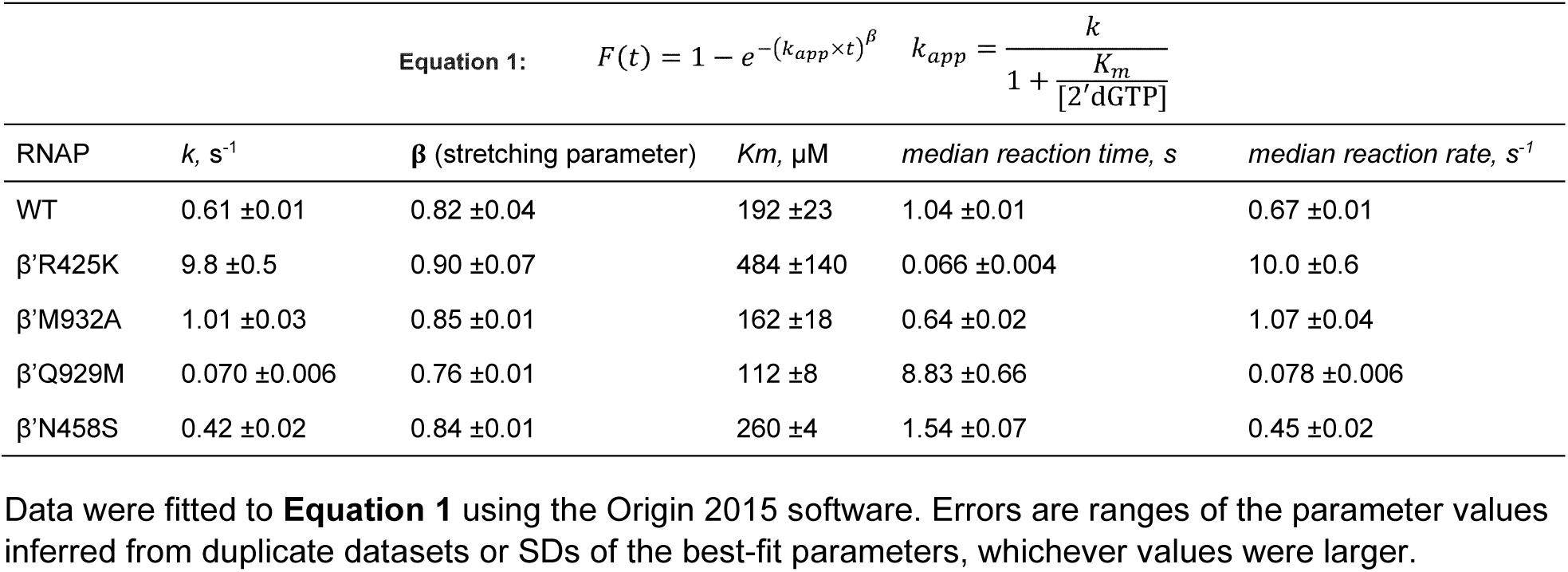
Kinetic parameters for the binding and incorporation of 2’dGMP by the wild-type and altered *E. coli* RNAPs.

**Supplementary Table 5.**
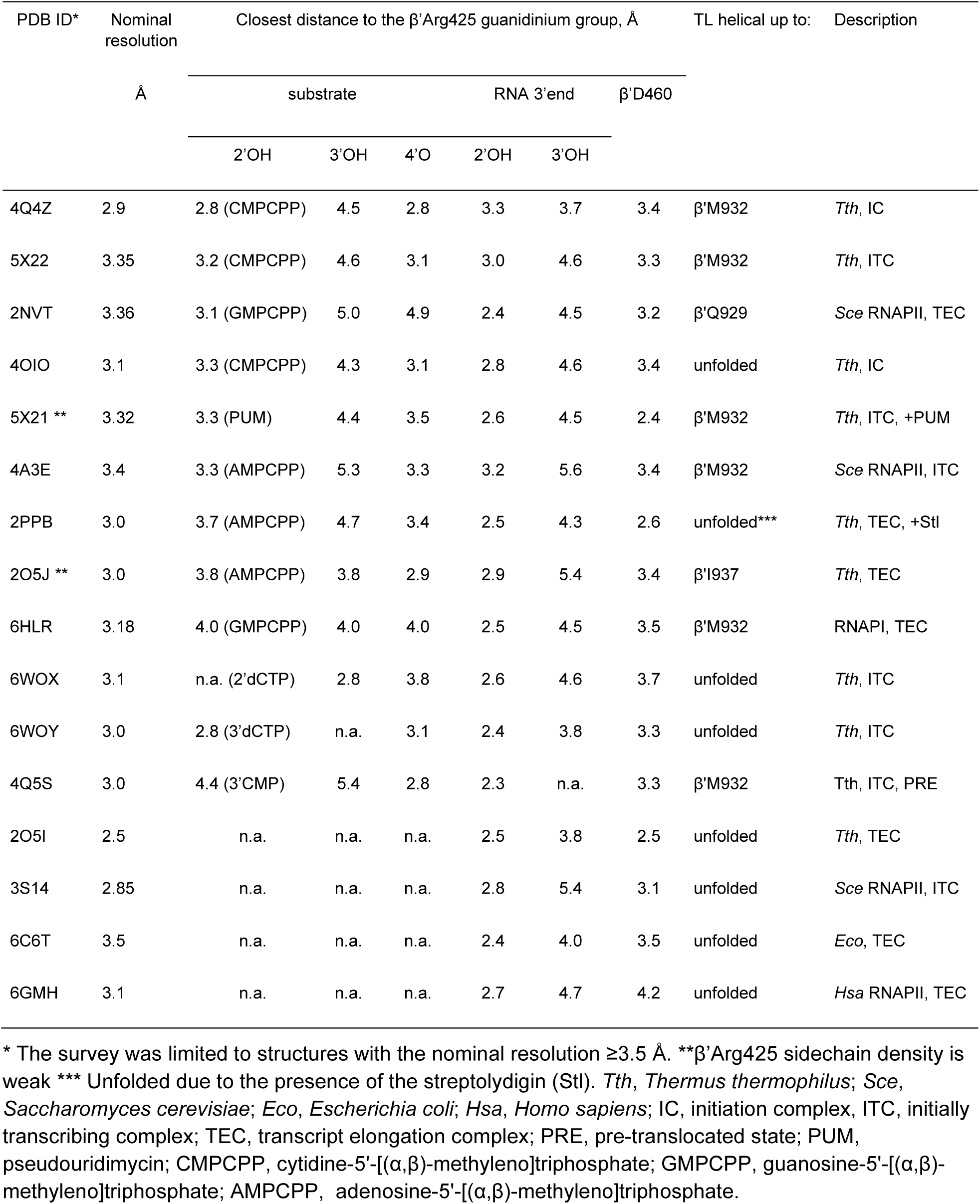
Distances between the guanidinium group of the β’Arg425 and the substrate, the RNA 3’ end and the β’D460 in structures of initiating, initially transcribing and elongating multi-subunit RNAPs.

**Supplementary Table 6.** The recovery frequency and affinities of the templated poses during docking of the 3’-endo CMP, 3’-endo 2’dCMP and 2’-endo 2’dCMP to the active site of *T. thermophilus* RNAP.

| Receptor | *Tth* IC<br>PDB ID 4Q4Z | *Tth* IC<br>PDB ID 4Q4Z | *Tth* IC<br>PDB ID 4Q4Z | *Tth* IC<br>PDB ID 4Q4Z | *Tth* ITC<br>PDB ID 821 | *Tth* ITC<br>PDB ID 821 |
| --- | --- | --- | --- | --- | --- | --- |
| Side chains | rigid | rigid | $\beta$'R425 (*Tth*<br>$\beta$'R704) flexible | $\beta$'R425 (*Tth*<br>$\beta$'R704) altered | rigid | rigid |
| TL | partially<br>folded | partially<br>folded | partially<br>folded | partially<br>folded | unfolded | unfolded |
| Original<br>ligand | 3'-endo<br>CMPCPP | 3'-endo<br>CMPCPP | 3'-endo<br>CMPCPP | 3'-endo<br>CMPCPP | 2'-endo<br>2'dCTP | 2'-endo<br>2'dCTP |
| Docked<br>ligand | 3'-endo CMP<br>PDB ID 3BSO | 3'-endo 2'dCMP<br>PDB ID 4O3N | 2'-endo 2'dCMP<br>PDB ID 3FL6 | 2'-endo 2'dCMP<br>PDB ID 3FL6 | 2'-endo 2'dCMP<br>PDB ID 3FL6 | 3'-endo 2'dCMP<br>PDB ID 4O3N |
| Run | Recovered templated poses and their affinity (kcal/mol) based on Autodock Vina empirical scoring function |  |  |  |  |  |
| 1 | -7.8, -7.7 | -6.9 | - | -7.1 | -6.8 | - |
| 2 | -7.8, -7.6 | - | - | -7.0, -6.9 | -6.6 | - |
| 3 | -7.9, -7.8 | -6.7 | - | -7.2, -7.0 | -6.6 | - |
| 4 | -7.8, -7.7 | -6.9, -6.5 | -6.9 | -7.0 | - | -6.4 |
| 5 | -7.8, -7.6 | - | -6.7 | -7.0 | -6.6 | - |
| 6 | -7.9, -7.7 | -6.8, -6.7 | - | -6.9 | -6.4 | - |
| 7 | -7.6, -7.5 | -6.3 | -7.1, -7.1 | -7.0 | -6.7 | - |
| 8 | -7.6, -7.5 | -7.1 | - | -7.0 | -6.7 | - |
| 9 | -7.9, -7.7 | -7.0 | -6.8 | -6.9 | -6.6 | -6.2 |
| 10 | -7.8, -7.6 | -6.8 | -6.6 | -7.1 | -6.7 | - |

## Supplementary Note

### Kinetic data analyses

We used time-resolved single nucleotide addition experiments to estimate the equilibrium constant for GTP, 2’dGTP and 3’dGTP binding and dissociation in the active site of RNAP and to determine the first order rate constant (also known as the turnover number) for the incorporation of GMP, 2’dGMP and 3’dGMP into the nascent RNA. The TECs were assembled on synthetic nucleic acid scaffolds and contained the fully complementary transcription bubble flanked by 20-nucleotide DNA duplexes upstream and downstream (**Supplementary Fig. 1a**). The annealing region of a 16-nucleotide RNA primer was initially 9 nucleotides, permitting the TEC extended by one nucleotide to adopt the post- and pre-translocated states, but disfavoring backtracking. The RNA primer was 5’ labeled with the infrared fluorophore ATTO680 to monitor the RNA extension by denaturing PAGE.

To facilitate the rapid acquisition of kinetic data (see below), the template DNA strand contained a fluorescent base analogue 6-methyl-isoxanthopterin (6-MI) eight nucleotides upstream from the RNA 3’ end. 6-MI allowed the monitoring of RNAP translocation along the DNA following nucleotide incorporation (**Supplementary Fig. 1a)**. 6-MI was initially positioned in the RNA:DNA hybrid eight nucleotides upstream of the RNA 3’end. The 6-MI fluorescence was quenched by the neighboring base pairs in the initial TEC and the pre-translocated TEC that formed following nucleotide incorporation, but increased when the 6-MI relocated to the edge of the RNA:DNA hybrid upon translocation. This fluorescence system was extensively validated in our previous studies ^28,29,56^.

We first measured concentration series of GMP and 2’dGMP incorporation by the wild-type and altered RNAPs using a time-resolved fluorescence assay performed in a stopped flow instrument (**Supplementary Figs. 1-3**). We used the translocation assay because it allowed the rapid acquisition of concentration series, whereas measurements of concentration series by monitoring RNA extension in a rapid chemical quench-flow setup would be considerably more laborious. We then performed a preliminary data analysis by fitting each fluorescence timetrace to a single exponential function followed by fitting the resulting individual rates to a Michaelis equation. The inferred *k*_*cat*_ and *Km* generally supported all major conclusions reported in this study. However, we proceeded to expand the datasets by including additional data and developed more elaborate analysis routines. The first reason to invoke a more elaborate analysis was the observation that most fluorescence time traces in our datasets fitted poorly to the single exponential function. In fact, the underlying physics of a single turnover enzymatic reaction suggests that individual timetraces in the concentration series should, in a general case, be poorly described by a single exponential function (see below). The second reason to invoke a more elaborate analysis was the concern that the Michaelis constant is a lumped constant that contains a sum of the catalytic and substrate dissociation rates in the numerator and the substrate binding rate in the denominator, whereas the equilibrium binding constants are the ratios of the substrate dissociation and binding rates. Accordingly, we were concerned that comparing the Michaelis constants of reactions could potentially lead to erroneous conclusions in the cases where the *Km* was markedly different from the *K*_*D*_.

For the sake of understanding our analysis workflow, it is important to acknowledge that each reaction timetrace in the concentration series describes a single turnover process: we designed the transcribed sequence so that only a single GMP (or 2’dGMP or 3’dGMP) became incorporated upon the addition of GTP (or 2’dGTP or 3’dGTP). The ease of obtaining single turnover timetraces is a significant analytical advantage natively associated with template-dependent nucleic acid polymerases. It is often possible to infer more parameters from concentration series of single-turnover reactions than from concentration series of classic multi-turnover enzymatic reactions.

Next, most timetraces in the concentration series are not expected to fit a single exponential function even in the case of the simple signal, a 1-nt extended nascent RNA (RNA17 in this study). The enzymatic reaction is minimally a two-step sequential reaction that consists of the substrate binding and the substrate incorporation steps. For timetraces obtained at sub-saturating [NTP], both reactions are partially rate limiting and the overall shape of the timetrace cannot be accurately described by a single exponential function. Only timetraces at saturating [NTP] (>10 × *Km*) and at the [NTP] far below saturation (<*Km*/10) are expected to fit well to a single exponential function (for RNA17). Furthermore, fluorescence timetraces do not fit the single exponential function even at the saturating [NTP], because the process leading to the change in the fluorescence signal consists of three steps: NTP binding, NMP incorporation and translocation. Both the NMP incorporation and translocation are partially rate limiting at saturating [NTP] ^29^ resulting in the time-traces not following a single exponential function. The considerations above indicate that the concentration dependencies should be fit globally to the two step (RNA17) or three step (fluorescence timetraces) sequential reaction scheme (**Scheme 1**) rather than be fit individually to a single exponential function.

We fitted the GTP, 3’dGTP and selected 2’dGTP concentration series to **Scheme 1** using the numerical integration capabilities of the Kintek Explorer software ^44^. The model postulated that the initial TEC16 reversibly binds the NTP substrate, undergoes an irreversible transition to TEC17 upon incorporation of the nucleotide into RNA, followed by an irreversible translocation. The concentration series data (fluorescence timetraces supplemented with a single HCl-quenched RNA extension curve at a saturating substrate concentration) allowed for the inference of the *k*_*cat*_, *K*_*D*_ and *k*_*tra*_ (in some cases), whereas the individual values of the *k*_*on*_ and *k*_*off*_ could not be resolved. Incorporation of an EDTA quench experiment allowed for the resolution of all parameters in **Scheme 1** in cases where the EDTA quenched curve was temporarily separated from the HCl quench curve (**Table 1**, GTP and 3’dGTP data; **Supplementary Table 3**). In cases where the EDTA quenched curve was not temporarily separated from the HCl quench curve, only the lower bounds of the *k*_*on*_ and *k*_*off*_ could be inferred (**Table 1**, 2d’GTP data). Overall, fits to **Scheme 1** were performed largely as described in Prajapati et al. ^45^ except that here we used a non-equilibrium heterogeneity model (see below) to describe the kinetic heterogeneity in the TEC preparations instead of a reversible inactivation model (see below) that was employed by Prajapati et al.

Next, a kinetic heterogeneity in the TEC preparations introduced an additional level of complexity to the fitting of the data. We reported previously that a vast majority of TECs contain 5-20% of a slow fraction that manifests itself as a slow phase in reaction timetraces of both the fluorescence signal (stopped-flow assay) and the extended RNA (quench flow assay) ^29,56^. In the case of fast reactions measured in this study (GTP, 3’dGTP data), the rates of the fast and slow phases differed approximately tenfold and therefore the phases could be precisely resolved (see a dedicated section below). Importantly, the fast phase of the reaction constituted 80-90% of the signal amplitude (**Table 1, Supplementary Table 3**). Accordingly, we considered the activity of the fast fraction as a representative measure of the RNAP activity in each experiment and disregarded the minor slow fraction when comparing the wild-type and variant RNAPs (**Fig. 2**).

In the case of slow reactions (2’dGTP data), the fast and the slow phases were not well separated (4-fold difference in rates, **Table 1**, 2’dGTP data for the wild-type RNAP). The poor separation led to large uncertainties in the *k*_*cat*_ and the percentage of the fast fraction. Moreover, the fitting algorithm partitioned the signal amplitude approximately equally between the fast and the slow phases so the activity of the fast fraction could not be used as a representative measure of the RNAP activity in each experiment (**Table 1**, 2’dGTP data for the wild-type RNAP). To circumvent this problem, we globally fit the 2’dGTP concentration series to a semi-empirical **Equation 1** instead of using **Scheme 1**.

When fitting data to **Equation 1**, each timetrace was described by a stretched exponential function (an empirical function that is often used to describe heterogeneous systems ^57^). At the same time, the exponent followed the hyperbolic dependence on the 2’dGTP concentration (**Supplementary Figs. 1c, 3**). Such fits described the data well and gave three parameters: a reaction rate constant (*k*), a stretching parameter (β) and the Michaelis constant (*Km*). When a stretching exponential function is applied to a process where the reactivity changes over time (or distance), the rate constant parameter (*k*) corresponds to the initial reaction rate constant. In our case, the stretched exponential fit potentially absorbed both temporal and structural heterogeneity as well as the deviations from the single exponential behavior caused by the sequential nature of the enzymatic reaction (see above). For this reason, the rate parameter (*k*) did not have an easily interpretable meaning. To circumvent this problem we calculated the median reaction time as (median reaction time) = (ln(2)^(1/β)) / *k*; then calculated the median reaction rate assuming that (median reaction rate) = ln(2) / (median reaction time) and used the median reaction rate as a measure when comparing the wild-type and variant RNAPs (**Fig. 2, Supplementary Table 4**). Next, fitting the data to **Equation 1** gives the *Km* rather than the *K*_*D*_. However, it is rather certain that *k*_*off*_ >> *k*_*cat*_ for all 2’dGMP incorporation reactions (**Table 1**, also see *Scenario 2* below). If so, *Km* approximately equals *K*_*D*_ for each 2’dGMP incorporation reaction. Accordingly, we used *Km* in place of *K*_*D*_ for 2’dGMP addition reactions when comparing substrates and RNAPs (**Fig. 2**).

Finally, we emphasize that the 2’dGMP incorporation data by the wild-type and the β’R425K RNAPs were fit to both **Scheme 1** and **Equation 1** leading to affinities for 2’dGTP that were indistinguishable within the margin of the experimental uncertainty (compare 2’dGTP data in **Table 1** and **Supplementary Table 4**). The catalytic activity of the wild-type RNAP towards 2’dGTP inferred by fitting the data to **Equation 1** was, as expected, in-between the catalytic activities of the fast and slow fraction inferred by fitting the data to **Scheme 1**. Accordingly, we argue that the employment of different analysis routines for GTP and 2’dGTP is of little concern for the main inferences drawn in this study.

### Handling of the translocation rate during the kinetic analysis of nucleotide binding and incorporation

We have previously shown that the nucleotide addition and the subsequent translocation along the DNA by the wild-type *E. coli* RNAP occur with similar rates at saturating concentrations of cognate NTPs ^29^. As a result, (*i*) the translocation timetraces are delayed by a few milliseconds relative to the nucleotide addition timecurves and (*ii*) the translocation timetraces at saturating concentrations of cognate NTP substrates are not well described by a single exponential function because both nucleotide addition and translocation are partially rate limiting. In this study, translocation rates were tangential to the main line of investigation, but they were necessary parameters during the global fitting of the fluorescence timetraces and GMP incorporation timecurves to **Scheme 1**. At the same time, the translocation rates are much faster than the 2’dGMP incorporation rates and could be completely disregarded during the analysis of the 2’dGTP concentration series by fitting the date to **Scheme 1** or **Equation 1**.

It should be noted that translocation rates reported in **Supplementary Table 3** should not be equated with the forward translocation rates. Thus, we modeled translocation as an irreversible transition in **Scheme 1**. As a result, the inferred translocation rates are the rates of the system approaching the translocation equilibrium after the nucleotide incorporation rather than the forward translocation rates. Albeit somewhat counterintuitively but following the rules of the formal kinetics the inferred equilibration rate equals the sum of the forward and the backward translocation rates. It was possible to further split the equilibration rate into the forward and backward translocation rates by assessing the completeness of the translocation, as we did in our previous studies ^56^. However, we refrained from doing so in this study because the translocation process was tangential to the main line of the investigation.

### Interpretation of EDTA quench experiments

EDTA inactivates the free NTPs by chelating Mg^2+^ but allows a fraction of the NTPs that are already bound in the RNAP active site to complete incorporation into the RNA. In contrast, HCl denatures RNAP so neither free nor RNAP-bound NTPs can be incorporated into the RNA after the addition of HCl. As a result, a comparison of the EDTA-quenched timecurve and HCl-quenched timecurve may provide information about the NTP dissociation rate from the active site. If the rate of NTP dissociation from the active site is comparable to (or smaller than) the catalytic rate, the EDTA-quenched curve reports more extended RNA than the HCl-quenched curve at each timepoint ^30^. In contrast, if the rate of NTP dissociation from the active site is much larger than the catalytic rate, the EDTA- and HCl-quenched curves superimpose. Consistently, Kireeva et al. showed that the EDTA quench experiment is fully equivalent to the pulse-chase setup when performed with the *S. cerevisiae* RNAPII ^31^. We explicitly modeled the EDTA quench experiments using the pulse-chase routine of the Kintek explorer software. Such an approach does not require a priori assumptions about the reaction rates in the three-step scheme employed for fitting the data. However, we consider it imperative to recognize the graphic signatures of the EDTA-quenched curves, rather than solely rely on the fitting algorithm as a “black box” to estimate the parameters. Accordingly, we discuss the shape of the EDTA-quenched curve under three scenarios with different ratios of *k*_*cat*_ and *k*_*off*_ and relate them to our data.

### Scenario 1. The rate of NTP dissociation from the active site (*k*_*off*_) is much lower than the rate of NMP incorporation (*k*_*cat*_)

In this case, EDTA quench curves of a fully post-translocated and kinetically homogeneous TEC are expected to fit a single exponential function with the exponent corresponding to the pseudo first order rate constant for NTP binding (*k*_*on*_ × [NTP]) (**Supplementary Note Fig. 1a**). As a result, both TEC17 and TEC16NTP are detected as TEC17 in the EDTA quenched samples because nearly 100% of the TEC16NTP is converted into TEC17 after the addition of EDTA, and practically no NTP dissociates back into the solution (*k*_*cat*_ >> *k*_*off*_). In this situation, EDTA quenched curves precede HCl quenched curves at saturating [NTP]: the HCl curve is limited by *k*_*cat*_, whereas the EDTA curve is limited by *k*_*on*_ × [NTP]. In this scenario, at least *k*_*cat*_ and *k*_*on*_ can be inferred from NTP concentration series alone because *Km* = (*k*_*cat*_ + *k*_*off*_)/*k*_*on*_ ≈ *k*_*cat*_/*k*_*on*_, so *k*_*on*_ ≈ *k*_*cat*_/*Km*. The global fit of the NTP concentration series and the EDTA quench data additionally allows the inference of the upper bounds of *k*_*off*_ and *K*_*D*_.

We did not encounter *Scenario 1* in this study though the β’M932A data are a borderline case that resembles *Scenario 1*. While both the upper and lower bounds of *k*_*off*_ could be determined (**Supplementary Table 3**) the inferred values were markedly smaller than the *k*_*cat*_ of 17 - 29 s^-1^ and the lower bound for *k*_*off*_ was therefore very diffuse: best fit 3.1 s^-1^, lower bound 0.8 s^-1^, upper bound 5.4 s^-1^.

### Scenario 2. The rate of NTP dissociation from the active site (*k*_*off*_) is much faster than the rate of NMP incorporation (*k*_*cat*_)

In this case, only TEC17 is detected as TEC17 in the EDTA quenched samples, because nearly 100% of the TEC16NTP loses its NTP after the addition of EDTA and practically no NMP gets incorporated into the RNA after the addition of EDTA (*k*_*cat*_ << *k*_*off*_). In this situation, EDTA quenched curves always superimpose with HCl quenched curves. At least *k*_*cat*_ and *K*_*D*_ can be estimated from the NTP concentration series alone because *Km* = (*k*_*cat*_ + *k*_*off*_)/*k*_*on*_ ≈ *k*_*off*_/*k*_*on*_ = *K*_*D*_. The global fit of the NTP concentration series and the EDTA quench data additionally allow the inference of the lower bounds of *k*_*on*_ and *k*_*off*_

The above situation corresponds to 2’dGMP addition by the wild-type and variant RNAPs. Fitting the 2’dGTP concentration series to a semi-empirical **Equation 1** allowed the estimation of *k*_*cat*_ and *Km*^*2’dGTP*^ ≈ *K*_*D*_^*2’dGTP*^ for the wild-type, β’R425K, β’M932A, β’Q929M and β’N458S RNAPs (**Supplementary Table 4**). For the β’R425K and the wild-type RNAP we additionally measured the EDTA quench curve, fitted the data globally to **Scheme 1** and inferred the lower bounds of *k*_*on*_ and *k*_*off*_. in addition to *K*_*D*_ (**Table 1, Supplementary Figs. 1c, 3a**).

### Scenario 3. The rate of NTP dissociation from the active site (*k*_*off*_) is similar to the rate of NMP incorporation (*k*_*cat*_)

In this case, EDTA quench curves of a fully post-translocated and kinetically homogeneous TEC obtained at saturating [NTP] are expected to fit a biexponential function with the first exponent corresponding to the pseudo first order rate constant of NTP binding (*k*_*on*_ × [NTP]) and the second exponent corresponding to *k*_*cat*_. For normalized data, the amplitude of the first exponent equals *k*_*cat*_/(*k*_*cat*_ + *k*_*off*_) and the amplitude of the second exponent equals *k*_*off*_/(*k*_*cat*_ + *k*_*off*_). In this situation, EDTA-quenched curves precede HCl-quenched curves at saturating [NTP]: The HCl curve is limited by *k*_*cat*_, whereas the fast phase of the EDTA curve is limited by *k*_*on*_ × [NTP]. As always, *k*_*cat*_ and *Km* can be inferred from the NTP concentration series, but neither *k*_*cat*_/*Km* ≈ *k*_*on*_ (as is in *Scenario 1*) nor *Km* ≈ *K*_*D*_ (as is in *Scenario 2*). In contrast, the global fit of the NTP concentration series and the EDTA quench data has the best resolving power in *Scenario 3*: *k*_*cat*_, *k*_*on*_, *k*_*off*_, and *k*_*tra*_ (in some cases) can be inferred from the data though the precision of the individual estimates varies greatly.

The above situation corresponds to the GMP addition by the wild-type and variant RNAPs (**Supplementary Table 3, Supplementary Figs. 1b, 2**) and the 3’dGMP addition by the wild-type RNAP (**Table 1, Supplementary Fig. 3c**). Only the wild-type RNAP data allowed for precise estimates of all parameters of **Scheme 1**. In the case of the β’R425K and β’Q929M variants, the EDTA- and HCl-quenched curves separated poorly resulting in diffuse upper bounds for *k*_*on*_ and *k*_*off*_. In the case of all variant RNAPs, the HCl-quenched curves and the fluorescence timetraces separated poorly resulting in diffuse upper bounds for *k*_*tra*_. Finally, the EDTA-quenched curve of the β’M932A RNAP featured little change in the signal (RNA17) within the measured time interval resulting in a diffuse lower bound for *k*_*off*_. With all that said, relatively precise estimates for *k*_*cat*_ and *K*_*D*_ were obtained for all RNAPs and used as measures for the comparison of the RNAP’s capabilities to bind and utilize various substrates (**Fig. 2**).

### Handling of the slow fraction during fitting to Scheme 1

The timecourses of the NMP incorporation by the wild-type *E. coli* TEC typically display a distinctive slow phase that represents 5-25% of the overall signal amplitude and features the rate of 0.1 - 3 s^-1^. In contrast, the major, fast phase of the reaction is approximately tenfold faster at saturating [NTP] (20 - 30 s^-1^ for GTP). The slow phase possibly represents an inactive TEC in equilibrium with the active TEC, a fraction of the TEC that slowly reacts with the NTP substrate or a combination of both. During the fitting of the data using the Kintek Explorer software, the slow phase can be modeled in two ways (**Supplementary Note Fig. 1b**). The first option is to invoke a reversible equilibrium between the active and inactive TEC and to introduce a virtual equilibration step prior to mixing of the TEC with the NTPs. We term this approach as the reversible inactivation model. The second option is to explicitly model the TEC preparation as two fractions that do not interconvert but incorporate NMP with different rates. The fractions of the slow and fast TEC are then allowed to vary as parameters during the fit. We term this approach as the non-equilibrium heterogeneity model.

The two models are largely indistinguishable if measurements are carried out at a single [NTP] and both models require two parameters to describe the slow phase: inactivation and recovery rates in the first case, and the slow fraction and its reaction rate in the second case (**Supplementary Note Fig. 1b**). However, the response of the slow phase to the decrease in the [NTP] differs between these two models. The reversible inactivation model predicts that the rate of the slow phase is independent of [NTP] and the slow phase is largely abolished as the [NTP] decreases. In contrast, the non-equilibrium heterogeneity model predicts that the rate of the slow phase decreases in unison with the rate of the fast phase as [NTP] decreases (both follow a hyperbolic dependence on [NTP]). In this study we analyzed all GMP and 3’dGMP incorporation datasets using the non-equilibrium heterogeneity approach to model the slow phase, because some datasets (e.g. β’Q929M, **Supplementary Fig. 2**) could not be adequately fit by the previously employed reversible inactivation model ^28,45,56^.

**Supplementary Note Fig. 1:**
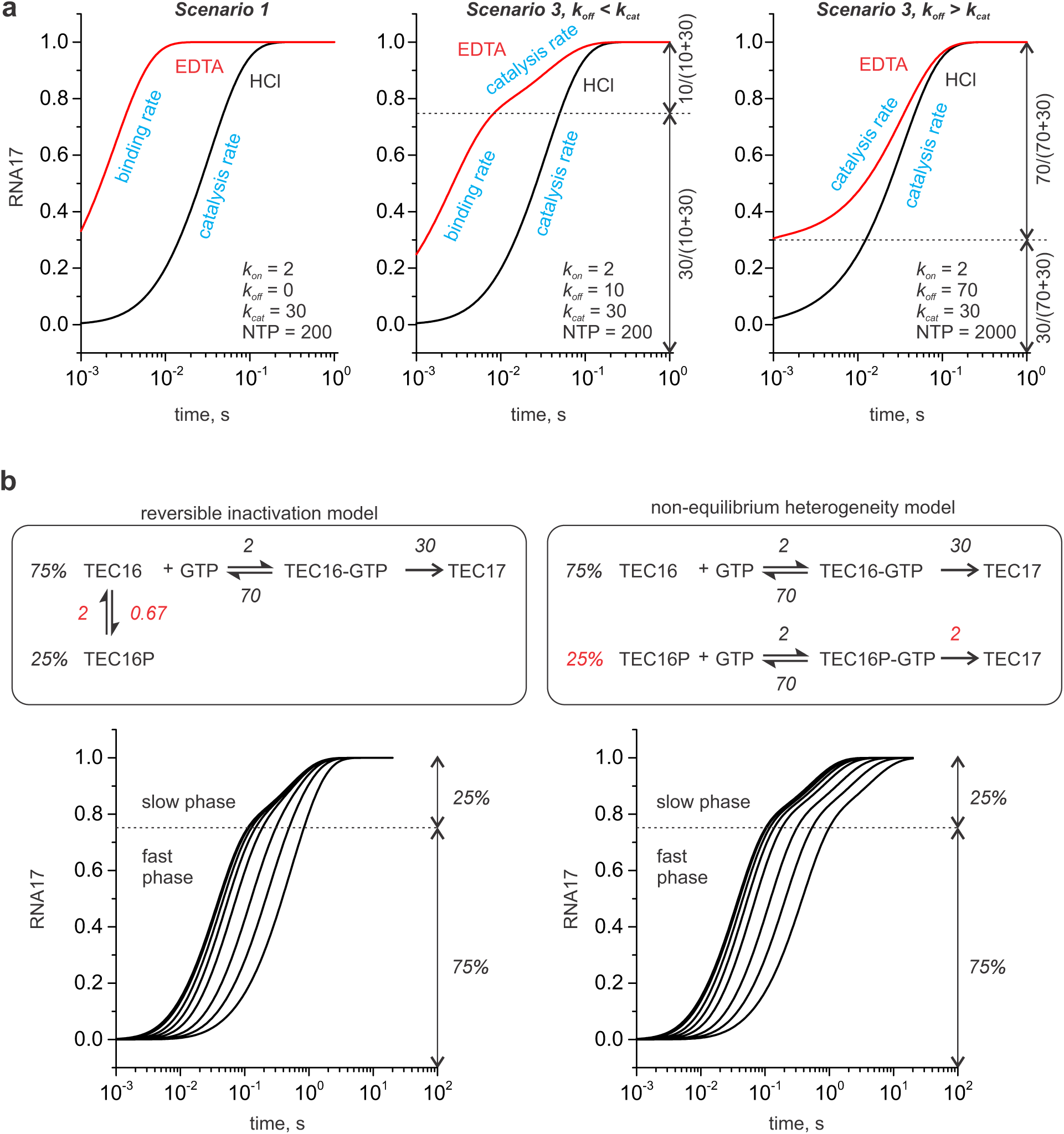
Kinetic analyses of the data. **a** Simulation and graphic interpretation of the EDTA and HCl quench curves at saturating substrate concentrations and different values of *k*_*off*_. **b** Simulation of concentration series of a biphasic reaction using the reversible inactivation (*left*) and non-equilibrium catalytic heterogeneity (*right*) models.

